# Enhancer Remodeling Promotes Tumor-Initiating Activity in NRF2-Activated Non-Small Cell Lung Cancers

**DOI:** 10.1101/2020.04.24.060798

**Authors:** Keito Okazaki, Hayato Anzawa, Zun Liu, Nao Ota, Hiroshi Kitamura, Yoshiaki Onodera, Md. Morshedul Alam, Daisuke Matsumaru, Takuma Suzuki, Fumiki Katsuoka, Shu Tadaka, Ikuko Motoike, Mika Watanabe, Akira Sakurada, Yoshinori Okada, Masayuki Yamamoto, Takashi Suzuki, Kengo Kinoshita, Hiroki Sekine, Hozumi Motohashi

**Author notes:** **Corresponding authors**, Hiroki Sekine, Ph.D., Department of Gene Expression Regulation, Institute of Development, Aging and Cancer, Tohoku University. 4-1 Seiryo-cho, Aoba-ku, Sendai, Miyagi, 980-8575, Japan. Hozumi Motohashi, M.D., Ph.D., Department of Gene Expression Regulation, Institute of Development, Aging and Cancer, Tohoku University. 4-1 Seiryo-cho, Aoba-ku, Sendai, Miyagi, 980-8575, Japan.

## Abstract

Transcriptional dysregulation, which can be caused by genetic and epigenetic alterations, is a fundamental feature of many cancers. A key cytoprotective transcriptional activator, NRF2, is often aberrantly activated in non-small cell lung cancers (NSCLCs) and supports both aggressive tumorigenesis and therapeutic resistance. Herein, we found that persistently activated NRF2 in NSCLCs generates enhancers at gene loci that are not normally regulated by transiently activated NRF2 under physiological conditions. Elevated accumulation of CEBPB in NRF2-activated NSCLCs was found to be one of the prerequisites for establishment of the unique NRF2-dependent enhancers, among which *NOTCH3* enhancer was shown to be critical for the promotion of tumor-initiating activity. Enhancer remodeling mediated by NRF2-CEBPB cooperativity promotes tumor-initiating activity and drives malignancy of NRF2-activated NSCLCs via establishment of the NRF2-NOTCH3 regulatory axis.

## Introduction

Transcriptional dysregulation is an important feature of many cancers (Bradner et al., 2017). Epigenetic alterations leading to aberrant transcription have been increasingly recognized in carcinogenesis ever since the emergence of the concept of the super-enhancer. These unique enhancer regions play a central role in the maintenance of cancer cell identity and drive oncogenic transcriptional programs to which cancer cells are highly addicted (Whyte et al., 2013; Lovén et al., 2013; Sengupta & George, 2017). Enhancer reprogramming, which is caused by the redistribution of transcription factors and the subsequent changes in transcription factor networks, drives cancer cell phenotypic drift during cancer initiation and progression (Chen et al., 2013; Denny et al., 2016). Thus, the elucidation of a cancer cell-specific enhancer landscape and transcription factors associated with the epigenetic environment is expected to provide powerful insights into the biological nature of cancer cells.

NRF2 (Nuclear Factor Erythroid 2 Like 2; NFE2L2) is a potent transcriptional activator that coordinately regulates many cytoprotective genes and plays a central role in defense mechanisms against oxidative and electrophilic insults (Yamamoto et al., 2018). Upon exposure to oxidative stress or electrophiles, NRF2 escapes KEAP1-mediated degradation and activates transcription through antioxidant response elements (AREs). While increased NRF2 activity is principally beneficial for our health, a variety of incurable cancers exploit NRF2 to achieve aggressive proliferation, tumorigenesis, and therapeutic resistance. Indeed, increased NRF2 accumulation in cancer tissues is strongly correlated with poor clinical outcomes in various cancer types (Shibata et al., 2008; Inoue et al., 2012; Onodera et al., 2014; Kanamori et al., 2015).

Persistent activation of NRF2 in cancer cells confers multiple advantages (Rojo de la Vega et al., 2018), such as increased survival due to enhanced antioxidant and detoxification capacities (Wang et al., 2008; Singh et al., 2010), increased proliferation as a result of metabolic reprogramming (Mitsuishi et al., 2012; Singh et al., 2013; DeNicola et al., 2015; Romero et al., 2017), protection of translational machinery from oxidative damage (Chio et al., 2016), and aggressive tumorigenesis resulting from the modulation of secretory phenotypes (Kitamura et al., 2017). In particular, NRF2 mediates drug resistance by increasing the expression of many detoxification enzymes and drug transporters (Maher et al., 2007; Hong et al., 2010), resulting in the inactivation and extrusion of small-molecule anti-cancer drugs. Due to these advantages, cancer cells with persistent NRF2 activation exhibit a heavy dependence on, or addiction to, NRF2 (Kitamura et al., 2018).

Therapeutic resistance is a major obstacle for the development of effective cancer treatments. Resistance may arise through genetic and/or epigenetic changes that are induced in cancer cells during treatment (Baylin, 2011). In particular, chemo- and radio-resistant tumor-initiating cells (TICs) impede treatment efficacy, thus leading to tumor relapse. Tumor-initiating abilities of cancer cells are experimentally evaluated based on their capacity to generate grossly recognizable tumors. Thus, the self-renewal capacity of TICs is not easily separated from their proliferative and survival abilities, which are strongly enhanced by NRF2, and chemo-resistant populations expressing high levels of NRF2 are often regarded as TICs (Mizuno et al., 2015; Jia et al., 2015; Ryoo et al., 2018). More precisely, it remains to be elucidated whether NRF2 does more than merely enhance proliferation and survival in order to support the tumor-initiating activity of cancer cells.

Approximately 15% of non-small cell lung cancer (NSCLC) cases carry somatic alterations of *KEAP1* gene, which are major causes of NRF2 addiction (Kandoth et al., 2013; Network, C.G.A.R. 2012; Weinstein et al., 2013; Network, C.G.A.R. 2014; Campbell et al., 2016). To clarify whether and how NRF2 contributes to the tumor-initiating activity and the consequent malignancy of NSCLCs exhibiting NRF2 addiction, we conducted an unbiased approach by investigating NRF2-dependent transcriptome in NSCLC cell lines with *KEAP1* mutations (NRF2-activated NSCLCs) and in those with an intact KEAP1-NRF2 system (NRF2-normal NSCLCs). We identified a battery of genes that are regulated by NRF2 specifically in NRF2-activated NSCLCs and found that these genes are accompanied by unique NRF2-dependent enhancers. CEBPB accumulation in NRF2-activated NSCLCs was found to be one of the prerequisites for the establishment of the unique enhancers, in which *NOTCH3* enhancer was critical for the promotion of tumor-initiating activity. Clinical data indeed showed that NOTCH3 contributes to cancer malignancy selectively in NRF2-activated NSCLCs, strongly suggesting the pathological significance of NRF2-NOTCH3 axis. The *NOTCH3* enhancer generated by NRF2 in cooperation with CEBPB establishes the NRF2-NOTCH3 axis and drives malignancy of NRF2-activated NSCLCs by promoting tumor-initiating activity.

## Results

### NRF2 promotes a stem-like phenotype of NRF2-activated NSCLCs

To clarify whether NRF2 has any active role in promoting tumor-initiating activity, which is one of the important properties for aggressive tumorigenesis (Extended Figures 1a and b), we cultured three NRF2-activated NSCLC cell lines with KEAP1 mutations, A549, H460 and H2023, under low attachment conditions in defined stem cell medium to allow them grow in the form of oncospheres (Justilien et al., 2014). TICs expressing stem cell markers were enriched in oncospheres growing under this condition (Extended Figure 1c). *NRF2* knockdown impaired oncosphere growth (Figures 1a-c), suggesting that NRF2 promotes a stem-like phenotype of NRF2-activated NSCLCs.

**Figure 1.**
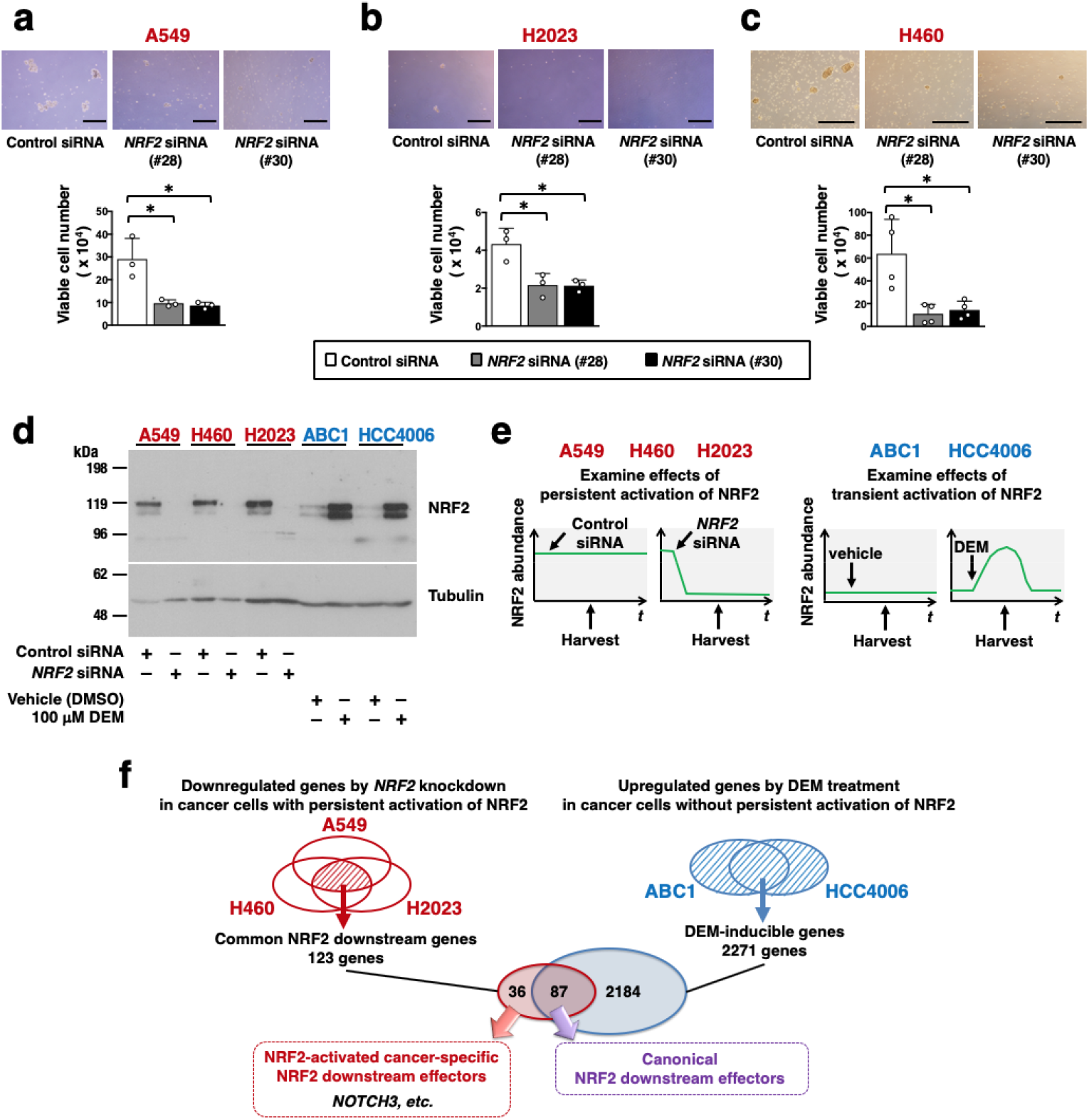
NRF2 enhances tumor-initiating activity in NRF2-activated NSCLC cell lines. **a-c.** Effects of *NRF2* knockdown on the oncosphere formation of A549 (**a**), H2023 (**b**) and H460 (**c**) cells. Scale bars indicate 100 μm, 50 μm and 500 μm, respectively (top panels). Viable cells were counted after trypsinization (bottom panels). Average cell numbers and SD from three independent experiments are shown. The Student’s *t* test was performed. \**p*<0.05. **d.** Immunoblot analysis of NRF2 protein levels in NSCLC cell lines. Tubulin was used as a loading control. One representative result of 3 independent experiments is shown. **e.** Experimental design for the preparation of RNA samples from NRF2-activated (left panels) and NRF2-normal (right panels) NSCLC cells. **f.** RNA-seq analysis of NSCLC cell lines. NRF2-activated (A549, H460 and H2023) and NRF2-normal (ABC1 and HCC4006) NSCLC cells were examined. In the following Figures, NRF2-activated and NRF2-normal NSCLC cells are indicated in red and blue, respectively.

### NOTCH3 is a unique downstream effector of NRF2 in NRF2-activated NSCLCs

Downstream effectors of NRF2 that promote tumor-initiating activity are potential therapeutic targets for suppressing tumorigenesis and cancer recurrence. We decided to identify such factors among NRF2 downstream effectors specific to NRF2-activated NSCLCs, so that their inhibition does not interfere with the cytoprotective functions of NRF2, which play beneficial roles in cancer-bearing hosts.

We first assessed NRF2 abundance in five NSCLC cell lines: three NRF2-activated (A549, H460 and H2023) and two NRF2-normal (ABC1 and HCC4006) NSCLC cell lines (Figure 1d). The NRF2-activated NSCLCs expressed high levels of NRF2, which could be diminished via transient transfection with an siRNA against NRF2. The two NRF2-normal NSCLCs with an intact KEAP1-NRF2 system expressed low levels of NRF2, which could be increased by treatment with an electrophile, diethylmaleate (DEM).

To identify unique downstream effectors of NRF2 in NRF2-activated NSCLCs, we compared NRF2-dependent transcriptomes across the three NRF2-activated NSCLC cell lines and the two NRF2-normal NSCLC cell lines. In the NRF2-activated cell lines, NRF2 was knocked down using two alternative siRNAs (#28 and #30) in order to evaluate genes dependent on persistently activated NRF2 (Figure 1e, left panel). In the NRF2-normal cell lines, NRF2 was induced using DEM to evaluate genes dependent on transiently activated NRF2 (Figure 1e, right panel). 123 genes were downregulated by *NRF2* knockdown in all three NRF2-activated cell lines (Figure 1f), which were defined as common NRF2 downstream genes. 2271 genes were upregulated by DEM treatment in either of the NRF2-normal cell lines (Figure 1f), which were defined as DEM-inducible genes. 87 genes that were included in both the common NRF2 downstream gene set and the DEM-inducible gene set were regarded as canonical NRF2 downstream effectors (Figure 1f). 36 genes included in the common NRF2 downstream gene set but not in the DEM-inducible gene set were regarded as NRF2-activated NSCLC-specific NRF2 downstream effectors (Figure 1f).

As a candidate for a regulator of a stem-like phenotype in NRF2-activated NSCLC cells, we selected NOTCH3 among the NRF2-activated NSCLC-specific NRF2 downstream effectors (Figure 1f and Extended Figure 2) based on a previous report showing that NOTCH3 contributes to a stem-like phenotype of NSCLC (Zheng et al., 2013; Ma et al., 2016).

### Co-expression of NRF2 and NOTCH3 is associated with a poor prognosis in lung adenocarcinoma cases

To examine whether the NRF2-NOTCH3 regulatory axis, which emerged in the cell line analysis, was also observed in clinical cases, we analyzed transcriptomic data from lung adenocarcinoma (LUAD) patients in the TCGA database (TCGA, provisional). The LUAD cases were divided into two groups according to their expression of representative NRF2 target genes: *NQO1, GCLC, GCLM, SLC7A11, TXNRD1* and *NR0B1. NOTCH3* expression was significantly higher in the LUAD cases with high NRF2 activity (Figure 2a). This result is consistent with our observation that *NOTCH3* is one of the NRF2-downstream effector genes unique to NRF2-activated NSCLC cell lines.

**Figure 2.**
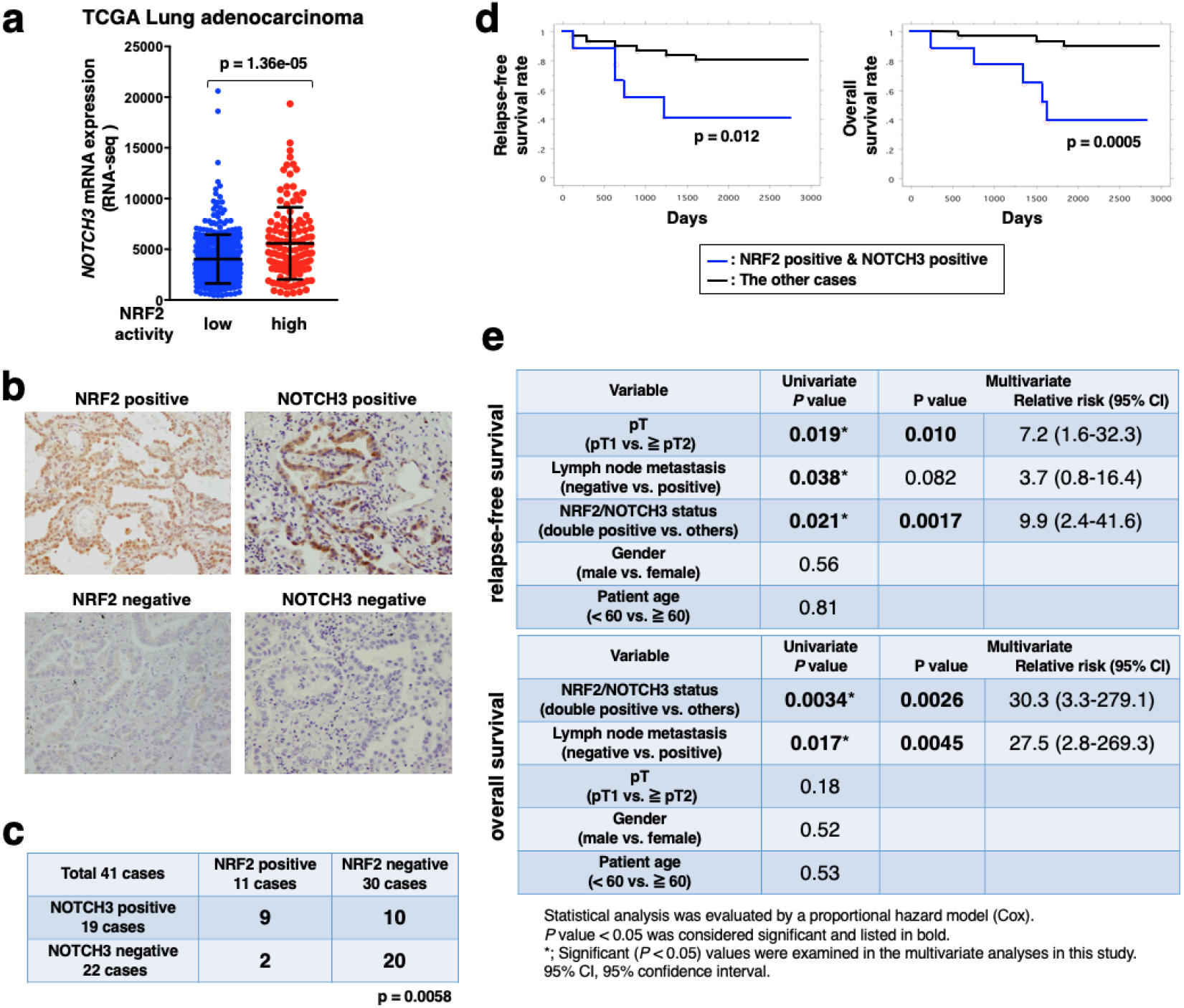
The NRF2-NOTCH3 axis is associated with poor clinical outcomes in lung adenocarcinoma patients. **a.** Comparison of *NOTCH3* mRNA expression in tumor tissues with low and high NRF2 activities. Gene expression data for lung adenocarcinoma patients were obtained from the TCGA database. The Student’s *t* test was performed. **b.** Immunostaining for NRF2 and NOTCH3 in human lung adenocarcinoma tissues. Representative cases with high and low expressions of NRF2 and NOTCH3 are shown. **c.** Association between NRF2 and NOTCH3 status in 41 lung adenocarcinoma patient samples. The Chi-square test was conducted to determine statistical significance. **d.** Overall survival rates and relapse-free survival rates for post-surgery patients grouped into NRF2 and NOTCH3 double-positive cases and the remaining cases. Kaplan-Meier analysis was conducted. Statistical significance was evaluated using the log-rank test. **e.** Univariate and multivariate analyses of 41 lung adenocarcinoma patients. The Cox proportional hazards model was used.

We next conducted a histological analysis of 41 LUAD samples from patients who had undergone surgical resection to examine a correlation of NRF2 and NOTCH3 statuses. The resected tumor tissues were stained with antibodies against NRF2 and NOTCH3 (Figure 2b). Most of the NRF2-positive cases, which are regarded as NRF2-activated cancers, were NOTCH3-positive (Figure 2c), which was again consistent with our observation that *NOTCH3* is a downstream effector of NRF2 in NRF2-activated NSCLC cell lines. In contrast, NOTCH3-positive cases were not necessarily NRF2-positive (Figure 2c), which suggested the presence of alternative regulators for *NOTCH3*. Notably, double-positive patients showed significantly poorer prognoses than the remaining cases (Figures 2d, e and Extended Figure 3), suggesting that the combination of NRF2 and NOTCH3 makes cancers malignant and that NOTCH3 contributes to cancer malignancy when it is co-expressed with NRF2 target genes in NRF2-activated NSCLCs. Although NOTCH3 has been reported to be associated with a poor prognosis of NSCLC (Ye et al., 2013), it appears to be the NRF2-NOTCH3 axis rather than NOTCH3 itself that contributes to the malignancy of NSCLCs.

### NRF2 is a direct activator of the *NOTCH3* enhancer

To decipher a mechanism connecting persistent activation of NRF2 and NOTCH3 expression, we examined whether NRF2 directly regulates the *NOTCH3* gene. We referred to a publicly available ChIP-seq data set of A549 cells in the ENCODE database. A clear NRF2 binding peak accompanied by its heterodimeric partner molecule MAFK was observed in the intergenic region between *NOTCH3* and *EPHX3* genes, approximately 10 kbp upstream of the *NOTCH3* transcription start site and 15-kb downstream of the *EPHX3* transcription termination site (Figure 3a). Three partially overlapping ARE sequences were present in the peak area.

**Figure 3.**
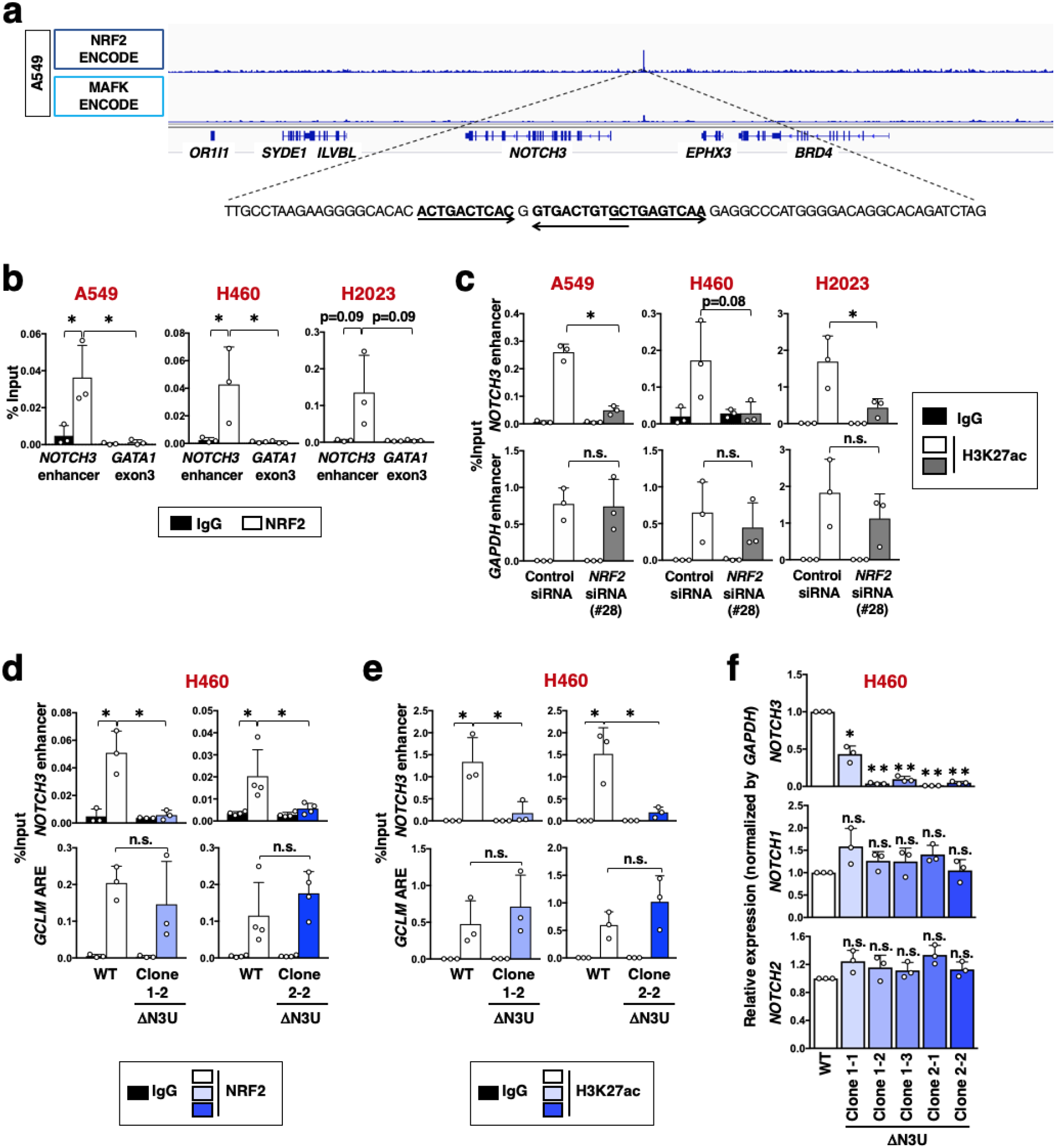
NRF2 directly activates *NOTCH3* expression. **a.** ChIP-seq profile of the *NOTCH3* locus in A549 cells. Data showing NRF2 and MAFK chromatin binding were obtained from the ENCODE database. **b.** ChIP assay using the NRF2 antibody in NRF2-activated NSCLC cells. Enrichment of the *NOTCH3* enhancer region was examined. *GATA1* exon 3 was selected as a control locus. The average and SD of three independent experiments are shown. The Student’s *t* test was performed. \**p*<0.05. **c.** ChIP assay using the H3K27ac antibody in three NRF2-activated NSCLC cells treated with control siRNA or *NRF2* siRNA. Enrichment of the *NOTCH3* enhancer region was examined. *GAPDH* enhancer was selected as a control locus. The average and SD of three independent experiments are shown. The Student’s *t* test was performed. \**p*<0.05, n.s.: not significant. **d, e.** ChIP assay using antibodies against NRF2 (**d**) and H3K27ac (**e**) in ΔN3U and wild-type (WT) H460 cells. Enrichment of the *NOTCH3* enhancer region was examined. The *GCLM* ARE was selected as a control locus. The average and SD of three or four independent experiments are shown. The Student’s *t* test was performed. \**p*<0.05, n.s.: not significant. **f.** RT-PCR measuring the expression levels of *NOTCH1, NOTCH2*, and *NOTCH3* normalized to *GAPDH* in ΔN3U H460 cells. Fold changes of the normalized values were calculated in comparison to WT H460 cells. The average and SD of the fold changes from three independent experiments are shown. Confidence interval estimation was conducted for the ΔN3U H460 clones. \**a*<0.05, \*\**a*<0.01, n.s.: not significant.

NRF2 binding at this site was clearly observed in all three NRF2-activated NSCLC cell lines (Figure 3b). Because NRF2 is known to recruit histone acetyltransferases CBP/p300 together with the SWI/SNF complex and Mediator complex (Katoh et al., 2001; Zhang et al., 2007; Sekine et al., 2015; Alam et al., 2017), NRF2-dependent deposition of acetylated histone H3K27 (H3K27ac), an enhancer mark, was also examined at this site. NRF2-dependent H3K27ac deposition was commonly observed in NRF2-activated NSCLC cell lines (Figure 3c), suggesting that this intergenic NRF2-dependent enhancer is a common feature of NRF2-activated NSCLCs.

To verify that this NRF2-dependent enhancer regulates *NOTCH3* expression, we disrupted the enhancer region in H460 cells by CRISPR-Cas9 genome editing technology (Extended Figures 4a). To exclude off-target effects of guide RNA (gRNA), two distinct gRNAs were used to obtain mutant clones (ΔN3U H460 cells) (Extended Figures 4a). NRF2 binding and H3K27ac deposition were abrogated in the mutated region in ΔN3U H460 cells (Figures 3d and e), verifying successful inactivation of the enhancer. *NOTCH3* expression was dramatically decreased (Figures 3f) but did not alter the expression of *EPHX3* or *BRD4* (Extended Figures 4b) in ΔN3U H460 cells, indicating that the intergenic NRF2 binding region serves as a major functional *NOTCH3* enhancer. No compensatory upregulation of *NOTCH1* or *NOTCH2* was observed (Figures 3f). In the mutant clones, NOTCH3 protein was hardly detected whereas NRF2 and its canonical target genes were not affected (Extended Figures 4c and d). The functional importance of this enhancer for *NOTCH3* expression was also verified in A549 and H2023 cells (Extended Figures 5a-d). Thus, NOTCH3 is one of the NRF2 target genes, being directly regulated by NRF2, in NRF2-activated NSCLCs.

### The *NOTCH3* enhancer is uniquely generated in NRF2-activated NSCLCs

Because RNA-seq analysis suggested that *NOTCH3* is an NRF2 target gene specifically in NRF2-activated NSCLCs, we further characterized *NOTCH3* expression and *NOTCH3* enhancer formation in relation to NRF2 activity in various cellular contexts.

*NRF2* knockdown decreased *NOTCH3* as well as *NQO1*, a canonical NRF2 target gene, in NRF2-activated NSCLC cells (Figure 4a). In contrast, physiological transient activation of NRF2 induced by DEM treatment elevated the canonical NRF2 targets *NQO1* and *GCLM* but not *NOTCH3* in NRF2-normal NSCLC cells (Figure 4b). These results are consistent with those obtained in the RNA-seq analysis, verifying that NRF2 does not regulate *NOTCH3* in NRF2-normal NSCLCs. Although a reciprocal relationship between NRF2 and the NOTCH pathway was previously reported (Wakabayashi et al., 2010; Wakabayashi et al., 2014), *NOTCH3* knockdown did not affect the expression levels of *NRF2* or *NQO1* in NRF2-activated NSCLC cells (Figure 4a).

**Figure 4.**
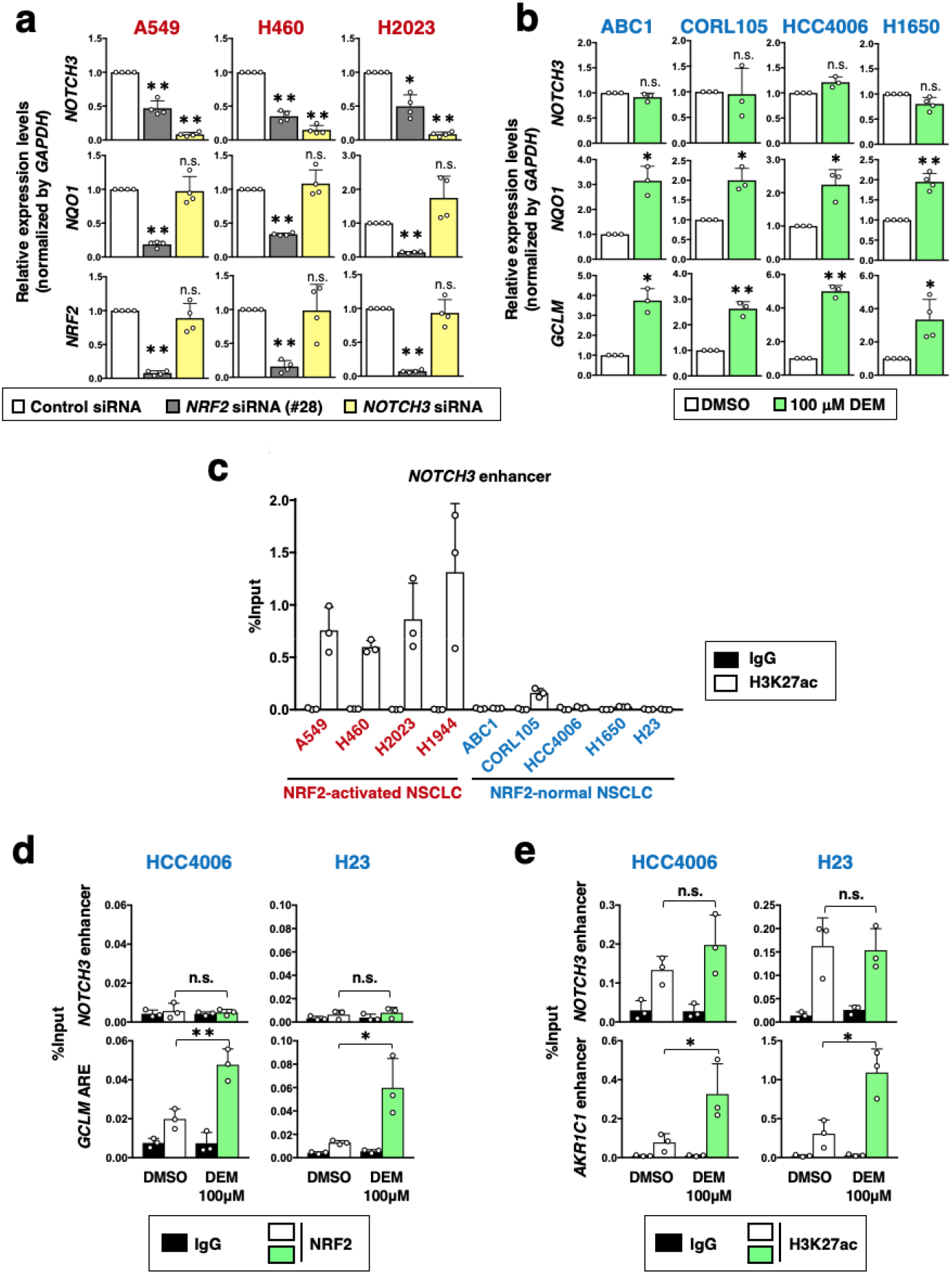
The *NOTCH3* enhancer is uniquely formed in NRF2-activated NSCLCs. **a.** RT-PCR measuring the expression of *NOTCH3, NQO1* and *NRF2* normalized to *GAPDH* in three NRF2-activated NSCLC cells treated with control siRNA, *NRF2* siRNA or *NOTCH3* siRNA. Fold changes of the normalized values were calculated in comparison to control siRNA treatment. The average and SD of the fold changes from four independent experiments are shown. Confidence interval estimation was conducted to evaluate statistical significance. \**a*<0.05, \*\**a*<0.01, n.s.: not significant. **b.** RT-PCR measuring the expression of *NOTCH3, NQO1* and *GCLM* normalized to *GAPDH* in four NRF2-normal NSCLC cells treated with vehicle (DMSO) or DEM. Fold changes **of** normalized values were calculated in comparison to vehicle-treated samples. The average and SD of the fold changes from three or four independent experiments are shown. Confidence interval estimation was conducted to evaluate statistical significance. \**a*<0.05, \*\**a*<0.01, n.s.: not significant. **c.** ChIP assay using the H3K27ac antibody in NRF2-activated and NRF2-normal NSCLC cells. The average and SD of 3 independent experiments are shown. **d, e.** ChIP assay using the NRF2 (**d**) or H3K27ac (**e**) antibodies in two NRF2-normal NSCLC cells treated with vehicle (DMSO) or DEM. The average and SD of 3 independent experiments are shown. The Student’s *t* test was performed. \**p*<0.05, \*\**p*<0.005, n.s.: not significant.

In mouse tissues, representatives of normal cells, pharmacological (CDDO-Im treatment) or genetic (*Keap1* knockdown) activation of NRF2 did not induce *Notch3*, whereas *Nqo1* was induced by both (Extended Figures 6a and b). These results suggest that NRF2 does not regulate *NOTCH3* in normal tissues. Thus, NOTCH3 was verified as a unique NRF2 target gene specifically in NRF2-activated NSCLCs.

In good agreement with this selective response of *NOTCH3* to NRF2 activation, the *NOTCH3* enhancer formation indicated by H3K27ac deposition was clearly detected in NRF2-activated NSCLCs but not in NRF2-normal NSCLCs (Figure 4c). Transient induction of NRF2 by DEM in NRF2-normal NSCLC cells, HCC4006 and H23, did not allow NRF2 binding or H3K27ac deposition at this site, whereas apparent increase was observed in canonical NRF2 target loci (Figures 4d and e). These results suggest that NRF2 binding and NRF2-mediated enhancer formation at the *NOTCH3* upstream region are restricted to NRF2-activated NSCLC cells.

### NRF2 generates a unique enhancer landscape in NRF2-activated NSCLCs

The highly selective attitude of NRF2 toward the *NOTCH3* enhancer attracted our interest in the enhancer landscape of NRF2-activated NSCLCs. We examined the genome-wide distribution of NRF2 and its contribution to enhancer formation by detecting H3K27ac deposition in A549 cells. We conducted ChIP-seq analysis of H3K27ac in A549 cells with or without *NRF2* knockdown and aligned the results with ENCODE NRF2 ChIP-seq data. NRF2 binding peaks were accompanied by H3K27ac deposition, which was reduced by *NRF2* knockdown (Extended Figure 7a and b), suggesting that NRF2 generally contributes to enhancer formation. If the log2 ratio of the H3K27ac levels in *NRF2*-knockdown cells versus control cells was less than –0.5, the H3K27ac deposition was defined as NRF2-dependent (Extended Figure 7c). Based on this definition, almost two-thirds of the H3K27ac deposition overlapping with NRF2-binding peaks were NRF2-dependent (Extended Figure 7c). A typical NRF2 binding motif was enriched in the NRF2-binding peaks that overlapped with NRF2-dependent H3K27ac deposition (Extended Figure 7d). Of 36 NRF2-activated NSCLC-specific NRF2 downstream effector genes, 21 genes, including *NOTCH3*, were regarded as direct NRF2 target genes based on the presence of NRF2 binding accompanied by the NRF2-dependent H3K27ac deposition (Figure 5a and Extended Figure 7e). Similarly, 77 out of 87 canonical NRF2 downstream effector genes were regarded as direct NRF2 target genes. The 77 canonical NRF2 target genes included well-known NRF2 target genes involved in cytoprotection and metabolism. Meanwhile, the 21 NRF2-activated NSCLC-specific NRF2 target genes were associated with a variety of biological functions.

**Figure 5.**
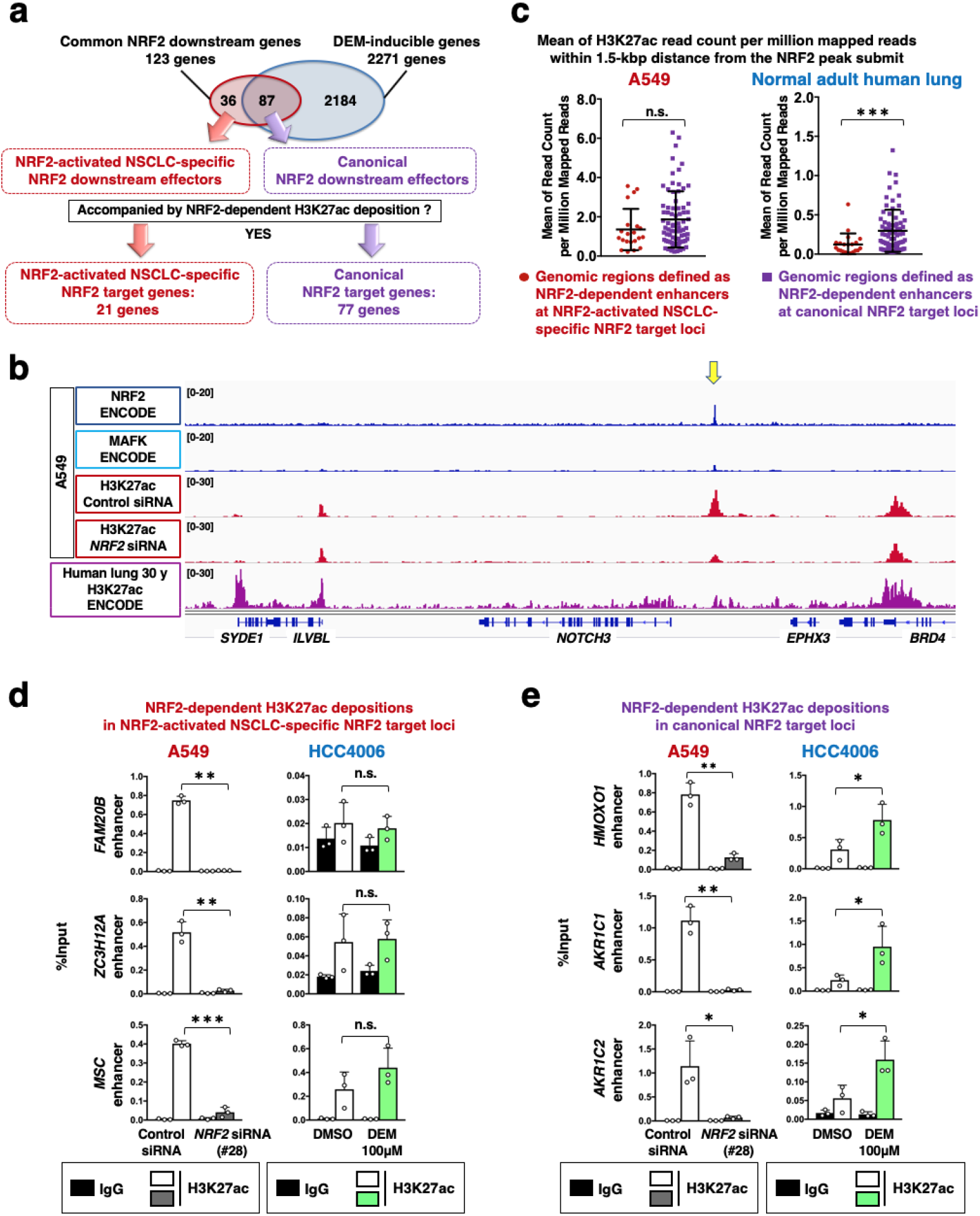
NRF2-activated NSCLC possesses unique enhancer signatures. **a.** NRF2-activated cancer-specific NRF2 target genes and canonical NRF2 target genes based on the presence of NRF2-dependent enhancers. **b.** ChIP-seq profiles at the *NOTCH3* locus. A549 cells (upper panels) and normal human lung samples (lower panel) are shown. NRF2 and MAFK chromatin binding in A549 cells and H3K27ac deposition patterns in normal human lung samples were obtained from the ENCODE database. Acetylated H3K27 deposition profiles in A549 cells treated with control siRNA or *NRF2* siRNA were obtained in this study. **c.** Comparison of H3K27ac deposition in NRF2-dependent enhancers that were defined in A549 cells. The NRF2-dependent enhancers were classified into 2 groups: those within NRF2-activated NSCLC-specific NRF2 target loci and those within canonical NRF2 target loci. H3K27ac deposition was compared between these 2 groups in A549 cells and normal adult lung samples. The Student’s *t* test was performed. \*\*\**p*<0.0005, n.s.: not significant. **d, e.** ChIP assay using the H3K27ac antibody in A549 cells treated with control or *NRF2* siRNA and HCC4006 cells treated with vehicle (DMSO) or DEM. H3K27ac deposition at NRF2-activated NSCLC-specific NRF2 target loci (**d**) and canonical NRF2 target loci (**e**) were examined. The average and SD of 3 independent experiments are shown. The Student’s *t* test was performed. \**p*<0.05, \*\**p*<0.005, \*\*\**p*<0.0005, n.s.: not significant.

Intriguingly, the majority of the H3K27ac depositions in NRF2-activated NSCLC-specific NRF2 target loci, including the one in the *NOTCH3* locus, were not clearly detected in normal adult human lung (Figure 5b, Extended Figures 8a and b). For instance, the one at *NR0B1* locus was strictly unique to NRF2-activated NSCLC cells (Extended Figure 8b), which is consistent with a previous study describing that NR0B1 is selectively expressed in *KEAP1*-mutant NSCLC cells (Bar-Peled et al., 2017). In contrast, H3K27ac depositions in the canonical NRF2 target gene loci were mostly observed both in A549 cells and normal adult human lung (Extended Figures 8c and d). More quantitatively, H3K27ac depositions at NRF2-activated NSCLC-specific NRF2 target loci were significantly lower than those at canonical NRF2 target gene loci in normal adult human lung, whereas in A549 cells, those in both loci were similarly high (Figure 5c).

We also compared the H3K27ac depositions in A549 and HCC4006 as representatives of NRF2-activated and NRF2-normal NSCLC cells, respectively. H3K27ac depositions in the NRF2-activated NSCLC-specific NRF2 target loci, *FAM20B, ZC3H12A* and *MSC*, were decreased by NRF2 knockdown in A549 cells but were not changed by DEM treatment in HCC4006 (Figure 5d). H3K27ac depositions in the canonical NRF2 target gene loci, *HMOX1, AKR1C1* and *AKR1C2*, were decreased by NRF2 knockdown in A549 cells and increased by DEM treatment in HCC4006 cells (Figure 5e).

Thus, H3K27 depositions at canonical NRF2 target gene loci are highly detectable in normal cells as well as in NSCLCs and enhanced by NRF2 activation. In contrast, those at NRF2-activated NSCLC-specific NRF2 target gene loci are uniquely observed in NRF2-activated NSCLCs and enhanced by NRF2 specifically in NRF2-activated NSCLCs. These results suggest that persistent NRF2 activation in cancer cells leads to the establishment of NRF2-dependent enhancers at gene loci that are not regulated by transiently activated NRF2.

### CEBPB colocalizes with NRF2 in active chromatin in NRF2-activated NSCLCs

To clarify a molecular mechanism by which NRF2 achieves the unique enhancer formation in NRF2-activated NSCLCs, we examined the genome-wide distributions of other transcription factors in A549 cells by utilizing ENCODE ChIP-seq data sets, anticipating that transcription factor cooperativity with NRF2 would be a key to understanding the establishment of unique NRF2-dependent enhancer signatures in the special cellular context, namely, NRF2-activated NSCLCs. Among 35 transcription factors with available ChIP-seq data of A549 cells in the basal condition, top 10 transcription factors colocalizing with NRF2 were selected (Figure 6a; Extended Table S1). MAFK ranked at the second place, which was reasonable as MAFK is a heterodimetic partner of NRF2. Although GATA3 and PBX3 ranked at the first and third place, respectively, their binding peaks were not robust at NRF2-dependent enhancers at 21 NRF2 target genes unique to NRF2-activated NSCLCs (data not shown). We thus chose CEBPB that ranked at the forth place for a further analysis. CEBPB colocalized with NRF2 in the active chromatin marked by H3K27ac deposition but not in the region without H3K27ac deposition (Figure 6b), implying a functional interaction between NRF2 and CEBPB. As expected, CEBPB was detected in the endogenous NRF2 transcription complex in A549 cells (Figure 6c). Notably, CEBPB protein levels in NRF2-activated NSCLC cells were higher than those in NRF2-normal NSCLC cells (Figure 6d), suggesting that a sufficient availability of CEBPB is one of the critical factors that invigorates NRF2 for the enhancer remodeling in NRF2-activated NSCLCs. Three out of six NRF2-dependennt enhancers unique to NRF2-activated NSCLCs, those in *FAM20B, ZC3H12A*, and *C5AR1* loci, exhibited reduced H3K27ac deposition by CEBPB knockdown (Figures 6e and f), suggesting that cooperativity between NRF2 and CEBPB explains a part of unique enhancer formation in NRF2-activated NSCLCs.

**Figure 6.**
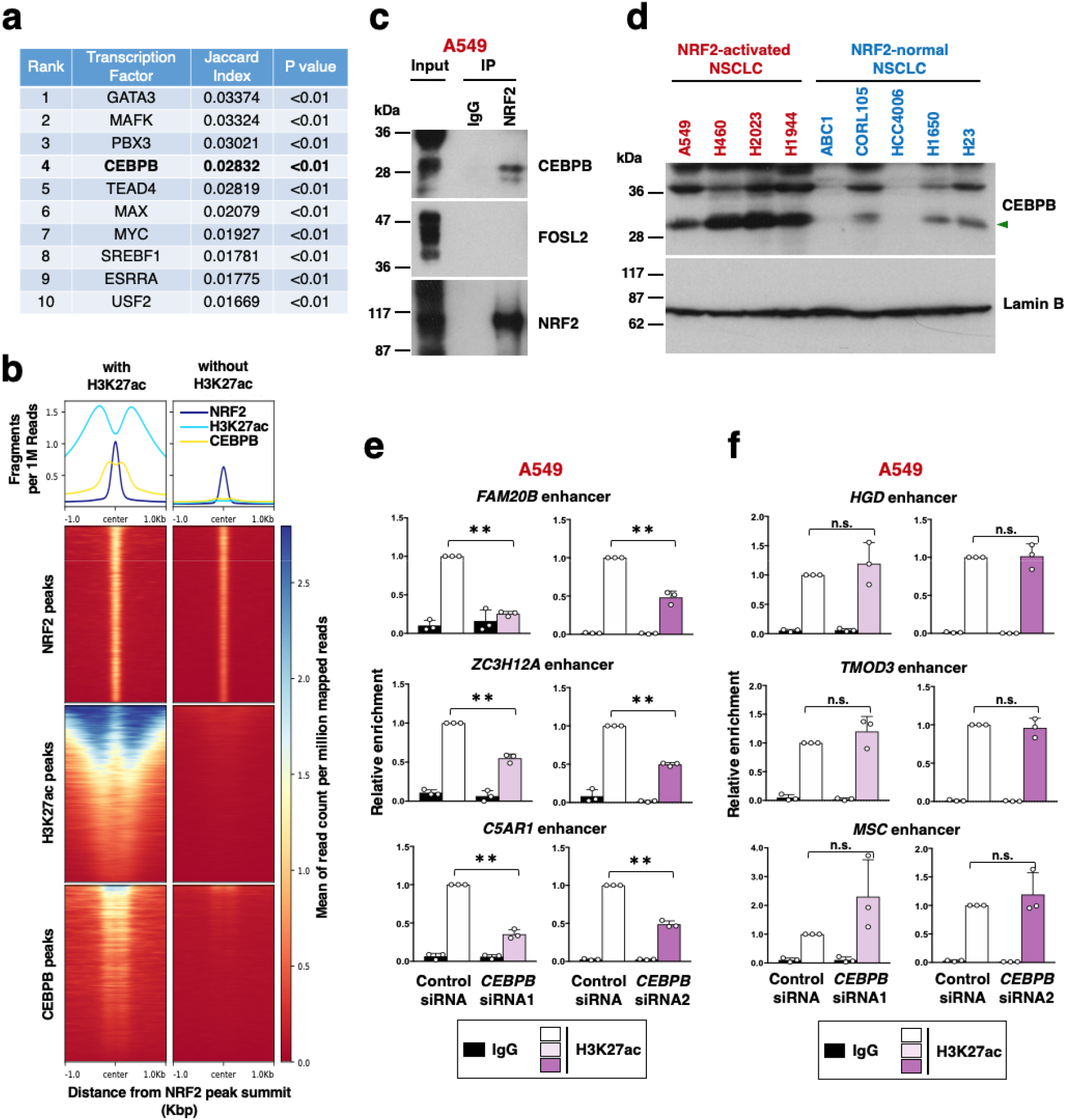
NRF2 cooperates with CEBPB for enhancer formation in NRF2-activated NSCLCs. **a.** Genome-wide colocalization of NRF2 and other transcription factors at active enhancers marked with H3K27ac deposition in A549 cells. Colocalization tendency was analyzed by Jaccard test using ChIP-seq data deposited in ENCODE. **b.** Aggregation plots and heat maps of ChIP-seq data of NRF2, H3K27ac and CEBPB surrounding NRF2 binding sites obtained from the ENCODE database. **c.** Affinity purification of the NRF2 complex from A549 cells. A representative result of 2 independent experiments is shown. **d.** Immunoblot analysis of CEBPB protein levels in nuclear extracts of four NRF2-activated and five NRF2-normal NSCLC cells. An arrowhead indicates the CEBPB isoform that corresponds to the one detected in the immunoprecipitation experiment with the NRF2 antibody shown in panel c. Lamin B expression was used as a loading control. The results shown are representative of two independent experiments. **e.** ChIP assay using the H3K27ac antibody in A549 cells treated with control or *CEBPB* siRNA. H3K27ac deposition at NRF2-activated NSCLC-specific NRF2 target loci were examined. The average and SD of 3 independent experiments are shown. The Student’s *t* test was performed. \*\**p*<0.005, n.s.: not significant.

### Cooperative chromatin binding of NRF2 and CEBPB generates *NOTCH3* enhancer in NRF2-activated NSCLCs

The *NOTCH3* enhancer region was found to be bound by seven transcription factors, including CEBPB, in A549 cells according to the ENCODE ChIP-seq data (Figure 7a) and contained consensus binding sites for eight transcription factors, among which those for NRF2/FOSL2 and CEBPB were conserved in the human and mouse (Figure 7b). Knockdown experiments in A549 cells revealed that CEBPB and FOSL2 made a substantial contribution to the *NOTCH3* expression (Figure 7c and Extended Figure 9a).

**Figure 7.**
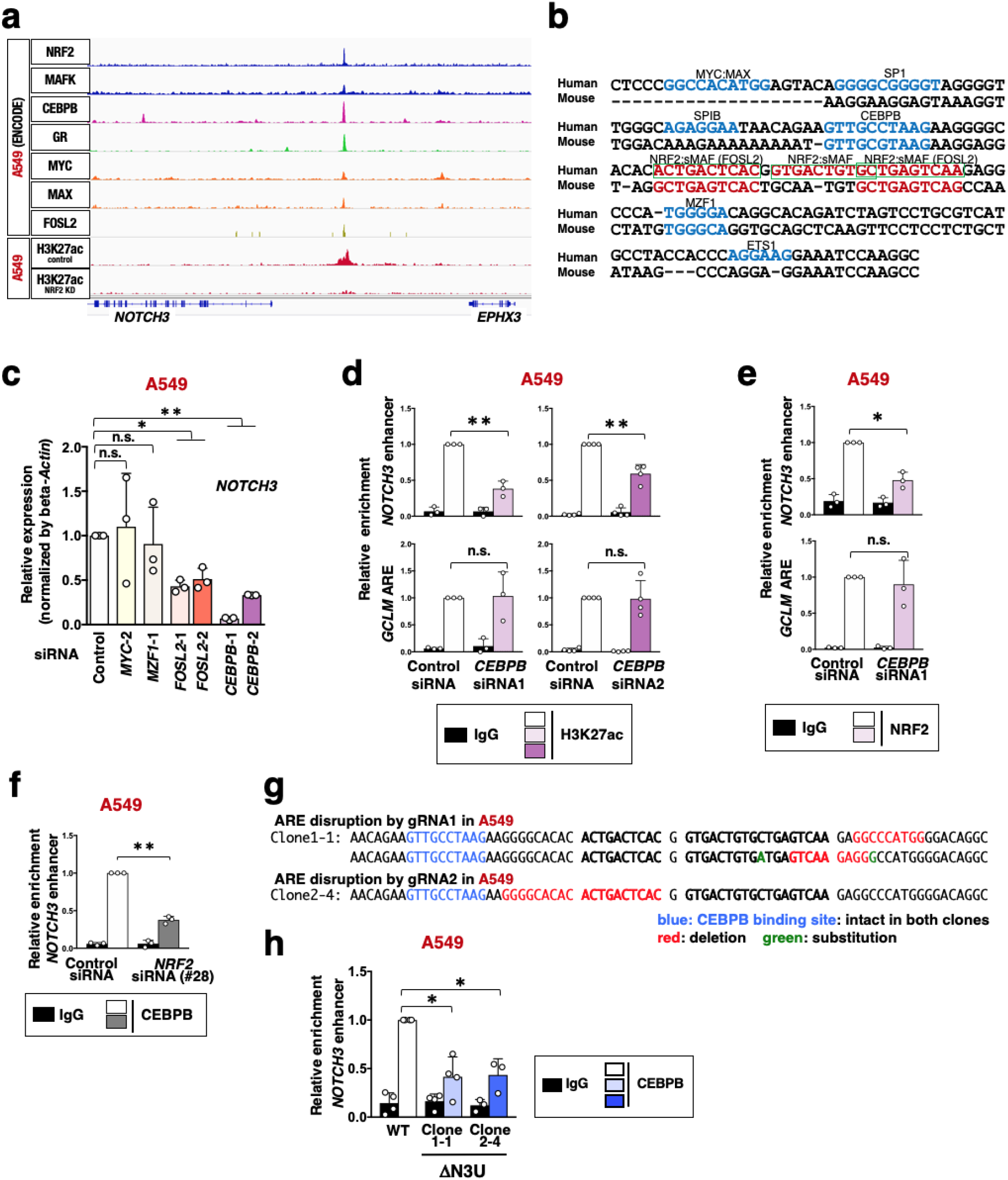
CEBPB-NRF2 cooperation is required for *NOTCH3* enhancer formation in NRF2-activated NSCLC cells. **a.** ChIP-seq profiles of the *NOTCH3* locus in A549 cells. **b.** Sequence comparison of flanking regions surrounding NRF2 binding sites in the *NOTCH3* enhancer between human and mouse. AREs (NRF2:sMAF binding sites) are shown in red, and three tandem human AREs are enclosed by green frames. Consensus binding motifs of other transcription factors are shown in blue. **c.** RT-PCR of *NOTCH3* in A549 cells treated with siRNAs against *MYC, MZF1, FOSL2, CEBPB* or control siRNA. Fold changes of the normalized values were calculated in comparison to A549 cells treated with control siRNA. The average and SD of the fold changes from 3 independent experiments are shown. Confidence interval estimation was conducted to evaluate statistical significance. \**a*<0.05, \*\**a*<0.01, n.s.: not significant. **d, e.** ChIP assay using the H3K27ac antibody (**d**) and NRF2 (**e**) in A549 cells treated with control siRNA or *CEBPB* siRNA. Enrichment of the *NOTCH3* enhancer region was examined. The *GCLM* ARE was selected as a control locus. Fold changes of %input values were calculated in comparison to the control samples incubated with H3K27ac or NRF2 antibody. The average and SD of 3 or 4 independent experiments are shown. Confidence interval estimation was conducted for knockdown samples incubated with H3K27ac or NRF2 antibody. \**a*<0.05, \*\**a*<0.01, n.s.: not significant. **f.** ChIP assay using the CEBPB antibody in A549 cells treated with control siRNA or *NRF2* siRNA. Fold changes of %input values were calculated in comparison to the control samples reacted with CEBPB antibody. The average and SD of 3 independent experiments are shown. Confidence interval estimation was conducted for knockdown samples reacted with H3K27ac antibody. \*\**a*<0.01. **g.** DNA sequence of ΔN3U A549 cells. A CEBPB binding site, shown in blue, is preserved in both ΔN3U A549 clones. Deletion and substitution by genome editing are shown in red and green, respectively. **h.** ChIP assay using the CEBPB antibody in ΔN3U and WT A549 cells. Fold changes of %input values were calculated in comparison to WT A549 cells incubated with CEBPB antibody. The average and SD of 3 or 4 independent experiments are shown. Confidence interval estimation was conducted for Clones 1-1 and 2-4 incubated with CEBPB antibody. \**a*<0.05.

At the *NOTCH3* enhancer region, *FOSL2* knockdown decreased H3K27ac deposition but did not affect NRF2 binding (Extended Figures 9b and c), and *NRF2* knockdown did not alter FOSL2 binding (Extended Figure 9d), indicating that FOSL2 contributes to *NOTCH3* enhancer formation independently of NRF2. In contrast, *CEBPB* knockdown decreased both H3K27ac deposition and NRF2 binding (Figures 7d and e), and *NRF2* knockdown decreased CEBPB binding (Figure 7f), suggesting the cooperative binding of NRF2 and CEBPB to the *NOTCH3* enhancer. Because CEBPB expression is decreased by NRF2 knockdown in A549 cells, we further verified the necessity of NRF2 for CEBPB binding to the *NOTCH3* enhancer by utilizing ΔN3U A549 cells, which harbor deletions in NRF2 binding sites (Figures 7g). In spite of the presence of an intact CEBPB binding motif, CEBPB binding to the *NOTCH3* enhancer was decreased in both clones of ΔN3U A549 cells (Figure 7h), verifying the cooperative binding of NRF2 and CEBPB.

A significant contribution of CEBPB to *NOTCH3* expression was also observed in other NRF2-activated NSCLC cells, H460 and H2023 (Extended Figures 9e and f), suggesting that NRF2-CEBPB cooperation is commonly important in NRF2-activated NSCLC cells. Thus, the *NOTCH3* enhancer comprises a unique enhancer landscape of NRF2-activated NSCLCs, which is shaped by NRF2-CEBPB cooperativity.

### The *NOTCH3* enhancer is critical for tumor-initiating activity of NRF2-activated NSCLCs

Finally, we examined the impact of the NRF2-NOTCH3 axis on the malignant behavior of NRF2-activated NSCLCs, which was suggested from the clinical study, in terms of tumor-initiating activity. Abrogation of the NRF2-NOTCH3 axis by disrupting the *NOTCH3* enhancer suppressed oncosphere growth of NRF2-activated NSCLC cell lines (Figure 8a, Extended Figures 10a and b). Knockdown of *CEBPB*, which was found to be a key cooperative factor with NRF2 for the *NOTCH3* enhancer formation, similarly suppressed the oncosphere growth of NRF2-activated NSCLC cell lines (Extended Figures 10c and d). Of note, disruption of the *NOTCH3* enhancer hardly affected spheroid growth (Figure 8b), which is a cell culture mode under low attachment conditions in normal media and reflects a simple cell growth ability in an anchorage-independent manner. These results are in a good contrast with those of *NRF2* knockdown in NRF2-activated cancers. *NRF2* knockdown impaired both oncosphere growth and spheroid growth (Extended Figure 10e and see Figures 1a-c), indicating that NRF2 promotes cell proliferation and survival in addition to tumor-initiating activity. Thus, the NRF2-NOTCH3 axis specifically contributes to tumor-initiating activity, which should be distinguished from other functional axes driven by NRF2 for malignant progression of cancers, such as cell proliferation and survival.

**Figure 8.**
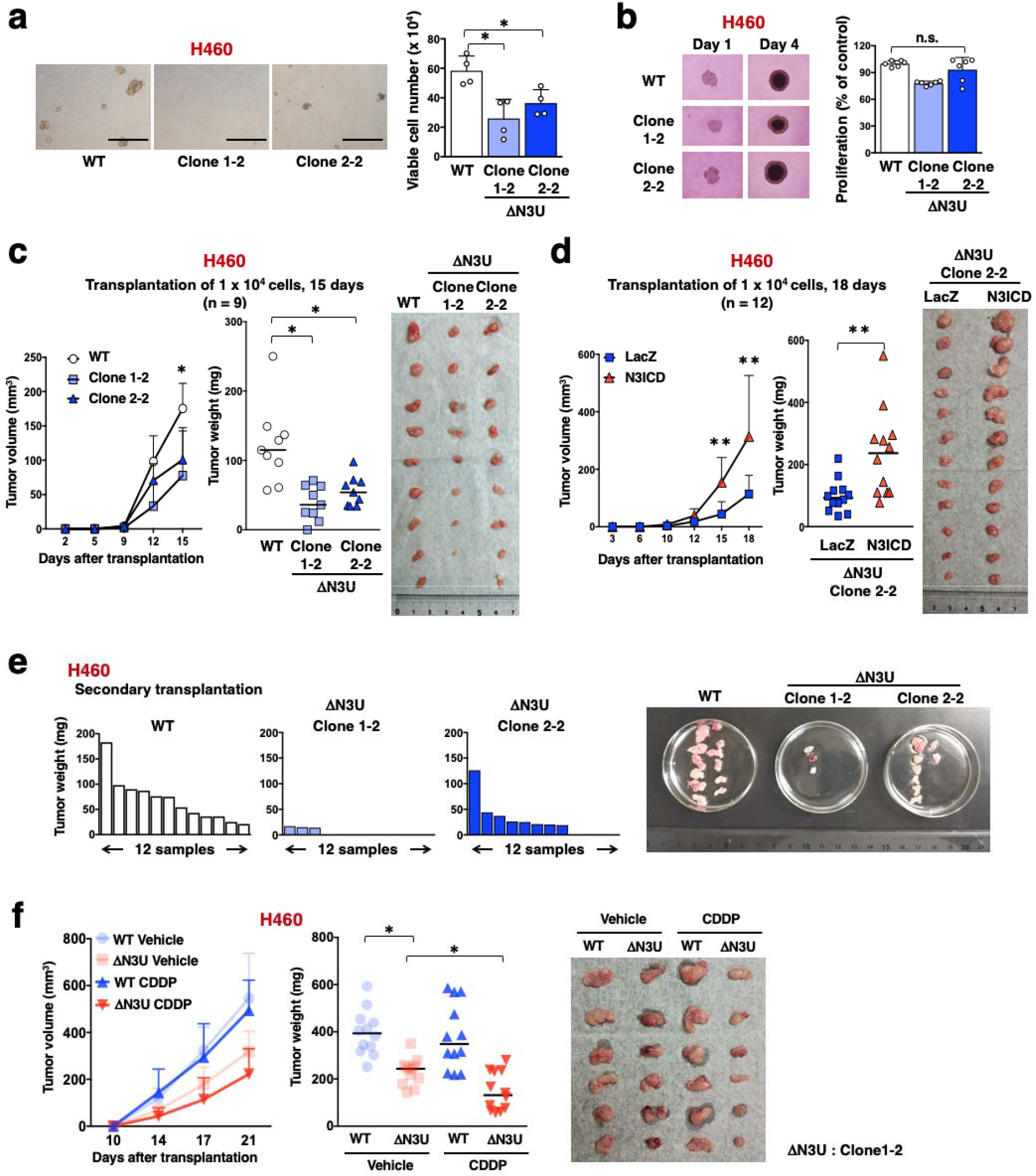
The *NOTCH3* enhancer promotes tumor-initiating activity of NRF2-activated NSCLCs. **a.** Oncosphere growth of ΔN3U and WT H460 cells (left panel). Scale bars indicate 500 μm. Viable cells were counted after trypsinization (right panel). Average cell numbers and SD from 4 independent experiments are shown. The Student’s *t* test was performed. \**p*<0.05. **b.** Spheroid growth of ΔN3U and WT H460 cells (left panels). Cell numbers were estimated using a cell counting kit on day 4 (right panel). Average cell numbers and SD from 6 independent experiments are shown. The average number of WT H460 cells was set as 100%. The Student’s *t* test was performed. n.s.: not significant. **c.** Xenograft experiment of ΔN3U and WT H460 cells. A photograph shows xenograft tumors at the time of tumor weight measurement on day 15. Horizontal bars indicate the median tumor weight. The Wilcoxon rank sum test was performed. \**p*<0.05. **d.** Xenograft experiment of ΔN3U H460 Clone 2-2 cells with LacZ and N3ICD. A photograph shows xenograft tumors at the time of tumor weight measurement on day 18. Horizontal bars indicate the median tumor weight. The Wilcoxon rank sum test was performed. \**p*<0.05, \*\**p*<0.005. **e.** Serial transplantation experiment of ΔN3U and WT H460 cells. Each graph shows the weight of tumors in the secondary transplantation. A photograph shows tumors at the time of weight measurement in the secondary transplantation. **f.** CDDP treatment experiment into nude mice after transplantation of ΔN3U and WT H460 cells. A representative photograph shows xenograft tumors at the time of tumor weight measurement on day 21. Horizontal bars indicate the median tumor weight. One-way ANOVA followed by the Bonferroni post hoc test was performed. \**p*<0.05.

We next conducted xenograft experiments to examine the *in vivo* contribution of the *NOTCH3* enhancer to the promotion of tumor-initiating activity in NRF2-activated NSCLC cells. The tumor growth of *NOTCH3* enhancer-disrupted cells, ΔN3U H460, ΔN3U A549 and ΔN3U H2023, was decreased compared with that of their parental cells (Figure 8c and Extended Figure 11). The suppressed tumor growth was restored by supplementation with the intracellular domain of NOTCH3 (N3ICD) (Figure 8d and Extended Figures 12), solidifying that the *NOTCH3* enhancer mediating the NRF2-NOTCH3 regulatory axis promotes tumor growth in NRF2-activated NSCLC cells. To further verify that the *NOTCH3* enhancer promotes the tumor-initiating activity of NRF2-activated NSCLC cells, we conducted a serial transplantation assay and compared the frequency of tumorigenesis (Figure 8e and Extended Figure 13). After the secondary transplantation, both clones of ΔN3U H460 cells generated a reduced number of tumors compared with parental H460 cells, supporting the notion that the NRF2-NOTCH3 axis contributes to the improved maintenance of tumor-initiating activity. Thus, *NOTCH3* enhancer inhibition targeting TICs was combined with cytotoxic anti-cancer drugs, such as cis-dichlorodiammineplatinum (CDDP), targeting the proliferating population for increased therapeutic efficacy. The tumorigenesis of ΔN3U H460 cells was suppressed by CDDP more effectively than that of parental cells (Figure 8f), successfully demonstrating a synergistic effect of CDDP and *NOTCH3* enhancer inhibition.

## Discussion

We linked persistent NRF2 activation in NSCLC cells to unique enhancer formation at the *NOTCH3* locus and demonstrated the clinical relevance of the NRF2-NOTCH3 regulatory axis. The significant anti-tumorigenic effect caused by the disruption of a single enhancer, *i.e*., disruption of the *NOTCH3* enhancer, stresses that enhancer dysregulation plays a critical role in the biological characteristics of cancers. The enhancer remodeling that occurs in NRF2-activated NSCLCs is partly mediated by a unique NRF2 transcription complex containing CEBPB. The accumulation of CEBPB is likely to modulate the NRF2 cistrome, which confers increased tumor-initiating activity on NRF2-activated NSCLCs by establishing the NRF2-NOTCH3 axis (Extended Figure 14).

NRF2 is capable of both inhibiting and promoting carcinogenesis depending on the cellular context (Taguchi et al., 2011). It is currently unclear both when and how NRF2 switches between its role as a guardian of cells that maintains redox homeostasis and its role as a driver of cancers that enhances aggressive tumorigenesis and therapeutic resistance. Based on studies demonstrating that sole overexpression of NRF2 in normal cells does not cause carcinogenesis, it is clear that constitutive activation of NRF2 by *KEAP1* or *NRF2* mutations, which are often encountered in NRF2-activated cancers, is not by itself a cancer driver (Wakabayashi et al., 2003; Taguchi et al., 2010; Noel et al., 2016; Suzuki et al., 2017; Murakami et al., 2017). In this study, we demonstrate that, in NRF2-activated NSCLC cells, NRF2 is involved in the transcriptional activation of non-canonical NRF2 target genes resulting from the enhancer dysregulation. We speculate that enhancer remodeling is required for the establishment of NRF2-activated cancers exhibiting NRF2 addiction. In this study, we found that CEBPB is one of the factors involved in the enhancer remodeling that invigorates NRF2 for transcriptional activation of non-canonical genes.

Among the various roles played by NRF2 in the promotion of malignant progression of cancers, the contributions of NRF2 to the maintenance of TICs were not fully understood. This study has shown that NRF2 promotes tumor-initiating activity in NSCLC cells by activating *NOTCH3*. This process seems to be distinct from other NRF2 functions such as increasing cell proliferation and survival, as disruption of the *NOTCH3* enhancer and the subsequent loss of *NOTCH3* expression had no effects on spheroid growth but impaired oncosphere growth of NRF2-activated NSCLC cells whereas *NRF2* knockdown impaired both. Our results strongly suggest that NRF2 contributes to the promotion of the tumor-initiating activity independently of its role in cell proliferation and survival by exploiting NOTCH3 as a unique downstream effector.

Our clinical study demonstrated that NRF2-NOTCH3 double-positive patients tended to have a significantly poorer prognosis compared with the others. These results suggest that not only NOTCH3 but other NRF2 canonical targets such as cytoprotective genes would also contribute to the poorer prognosis in NRF2-activated NSCLC. An important consideration here is that the promotion of tumor-initiating activity in TICs must be coupled with enhanced proliferation and survival of differentiated cancer cells in order to make a substantial contribution to cancer malignancy.

As a therapeutic perspective, NRF2 would be the best target for eliminating NRF2-activated cancers. Indeed, enthusiastic efforts are being made by many laboratories and pharmaceutical companies to develop NRF2 inhibitors for cancer cases that exhibit abnormal NRF2 activation and are therefore refractory to normal chemo- and radiotherapies (Ren et al., 2011; Singh et al., 2016; Tsuchida et al., 2017; Choi et al., 2017). However, considering the critical protective functions of NRF2, systemic administration of NRF2 inhibitors may cause deleterious effects in cancer-bearing hosts (Enomoto et al., 2001; Nezu et al., 2017). In this study, we explored unique NRF2 target genes and identified NOTCH3 as a key regulator of the tumor-initiating activity in NRF2-activated NSCLCs. Because *Notch3*-deficient mice exhibit modest vascular phenotypes but no serious defects (Kofler et al., 2015; Krebs et al., 2003) and because NOTCH3 is not involved in NRF2-mediated cytoprotection, NOTCH3 inhibition is expected to exert anti-cancer effects without interfering with normal cellular functions in cancer-bearing hosts. As NRF2-activated cancer cells are capable of highly efficient drug extrusion, targeting NOTCH3 is further advantageous in that it enables the circumvention of NRF2-mediated chemo-resistance. This is because NOTCH3 can be antagonized from the outside of cells at its functionally important extracellular domain. Indeed, an antagonizing antibody against NOTCH3 has potent anti-tumorigenic effects on xenografts of NRF2-activated NSCLC cell lines (unpublished observation). Therefore, NOTCH3 inhibition is expected to efficiently reduce the recurrence of NRF2-activated cancers by suppressing tumor-initiating activities without having adverse effects on cancer-bearing hosts.

Enhancer signatures have been shown to be more dynamic according to cell types and tissue types than those of promoters (Ernst et al., 2011; Roadmap Epigenomics Consortium et al., 2015; Nakato et al., 2019), which indicates that the same gene expressed in different cell types is often regulated by the same promoter but by different enhancers. Cancer-specific enhancers are expected to be ideal therapeutic targets because their inhibition would not interfere with the expression of respective genes in normal tissues. Modified nucleic acids and nucleic acid mimetics are possible candidates enabling the control of a specific enhancer by invading the DNA double strand and forming a triplex at the enhancer region of interest (Malnuit et al., 2011; Pabon-Martinez et al., 2017). Clarification of enhancer signatures responsible for the cancer malignancy and intervention to interfere with enhancer activities is an ambitious future challenge in development of anti-cancer drugs.

## Materials and Methods

### Cell culture

NRF2-activated NSCLC cell lines (A549, NCIH460 and NCIH2023 and NCIH1944) and NRF2-normal NSCLC cell lines (ABC1, CORL105, HCC4006, NCIH1650 and NCIH23) were used in this study. A549 cells were maintained in high glucose DMEM supplemented with 10% fetal bovine serum (FBS) and penicillin/streptomycin (Gibco). The rest of the cells were maintained in low glucose DMEM supplemented with 10% FBS and penicillin/streptomycin (Gibco). The cells were maintained in a 5% CO_2_ atmosphere at 37°C. PCR was used to confirm that the cultured cells were not infected with mycoplasma.

### Patients and tissue specimens

Tumor tissue specimens were obtained from 41 lung adenocarcinoma patients who underwent surgical resection between 2003 and 2004 in the Department of Thoracic Surgery at Tohoku University Hospital. The mean patient age was 66 years (range 37-82 years), with the exception of one case whose age was unknown. The mean follow-up period was 1982 days (range 233-2949). All specimens were fixed in 10% formalin and embedded in paraffin wax. All research protocols involved in this study were approved by the Ethics Committees at Tohoku University Graduate School of Medicine.

### Immunocompromised mice

Four-week-old male BALB/cAJcl-nu/nu mice (CLEA Japan) were used in this study. All animals were housed in specific pathogen-free conditions, according to the regulations of The Standards for *Human Care and Use of Laboratory Animals of Tohoku University* and the *Guidelines for Proper Conduct of Animal Experiments* by the Ministry of Education, Culture, Sports, Science, and Technology of Japan.

### Immunoblot analyses

For preparation of nuclear lysates, cells were lysed in buffer A (10mM HCl (pH7.5) 10mM KCl, 1.5mM MgCl_2_, 0.1 % NP40), and crude nuclei were pelleted by centrifugation and lysed in 2x Laemmli buffer followed by boiling at 95°C for 5 min. For preparation of whole-cell lysates, cells were directly lysed in 2x Laemmli buffer followed by boiling at 100°C for 10 min. The protein samples were separated by SDS-PAGE and transferred onto PVDF membranes (Immobilon P, Millipore). The antibodies used are as follows: anti-NRF2 (sc-13032X, Santa Cruz), anti-NOTCH3 (ab23426, Abcam), anti-CEBPB antibody (sc-150 X, Santa Cruz), anti-Tubulin (T9026, Sigma) and anti-Lamin B (sc-6217, Santa Cruz).

### Transient knockdown experiments

*NRF2* siRNAs were purchased from Invitrogen (cat. no. HSS107128 and HSS107130). *CEBPB* siRNAs (SASI_Hs01_00236022 and SASI_Hs01_00236027), *FOSL2* siRNAs (SASI_Hs01_00057657 and SASI_Hs02_00339278), *MYC* siRNA (SASI_Hs01_00222676) and *MZF1* siRNA (SASI_Hs01_00096728) were purchased from Sigma-Aldrich. MISSION siRNA Universal Negative Controls (Sigma-Aldrich) or DS scrambled negative control siRNA (IDT) were used as controls. siRNAs were transfected into cells either by electroporation using an MP-100 MicroPorator (Digital Bio Technology) or by lipofection using Lipofectamine™RNAiMAX Transfection Reagent (Thermo Fisher Scientific). Culture media were changed 24 hrs after transfection. After another 24-48 hrs, the cells were harvested for RNA purification, immunoblot analysis and the ChIP assay. The protocols used for transient knockdown with spheroid formation assay and oncosphere formation assays in each cell line are described below.

### Transient induction of NRF2

Cells were treated with 100 μM of diethyl maleate (DEM) or 30 nM of CDDO-Im and dimethyl sulfoxide (DMSO) as a vehicle. Cells were harvested either 4 hrs after treatment for immunoblot analysis and the ChIP assay or 16 hrs after treatment for RNA purification.

### RNA purification and RT-PCR

Total RNA was prepared from cells using ISOGEN (NIPPON GENE). cDNA was synthesized from 0.5 μg of total RNA using ReverTra Ace qPCR RT Master Mix with gDNA Remover (TOYOBO). Quantitative real-time PCR was performed for each sample in duplicate with Probe qPCR Mix (TOYOBO), SYBR qPCR Mix (TOYOBO) or PowerUp SYBR Green Master Mix (Thermo Fisher Scientific) using the Applied Biosystems 7300 PCR system (Applied Biosystems) and Applied Biosystems Quant Studio 3 (Applied Biosystems). The sequences of all primers used are listed in Extended Table S2. *GAPDH, HPRT* and *ACTB* were used as internal controls for normalization.

### RNA-sequencing analysis

Total RNA was extracted from A549, H460 and H2023 cells 48 hrs following transfection with control siRNA or NRF2 siRNA (cat. no. HSS107128 and HSS107130) using the RNeasy Mini Kit (Qiagen). Cells treated with the control siRNA were analyzed in biological duplicates. The cells treated with two different NRF2 siRNAs (HSS107128 and HSS107130) were analyzed separately and were considered to be biological duplicates of the NRF2-knockdown sample. Total RNA was extracted from ABC1 and HCC4006 cells 16 hrs after treatment with 100 μM DEM or vehicle (DMSO). Cells treated with either 100 μM DEM or vehicle (DMSO) were analyzed in biological duplicates. 4 μg of total RNA from A549 cells was subjected to rRNA removal using the Ribo-Zero Gold kit (Illumina). cDNA sequencing libraries were then prepared using the SureSelect Strand-Specific RNA library preparation kit (Agilent Technologies) with a modified protocol omitting the polyA selection step. Total RNA from all other cell lines was used to prepare cDNA sequencing libraries using the SureSelect Strand-Specific RNA library preparation kit (Agilent Technologies) after the polyA selection step. The libraries were quantified by qMiSeq and sequenced on a NextSeq 500 (Illumina) to generate 80-base paired-end reads. Data analysis was performed on the Illumina BaseSpace platform (https://basespace.illumina.com) as follows. Raw fastq sequencing files were aligned to the hg19 reference genome using TopHat Alignment Version 1.0.0 (Trapnell, et al., 2010). After read mapping, transcripts were assembled using Cufflinks software Version 2.3.1 (Trapnell, et al., 2010). Expression level estimations were reported for each sample as fragments per kilobase of transcript sequence per million mapped fragments (FPKM). Raw and processed data were deposited in GEO (GSE118840, GSE118842). To identify NRF2 downstream effectors, significantly decreased genes by *NRF2* knockdown were selected for A549, H460 and H2023 cells, and significantly increased genes by DEM treatment were selected for ABC1 and HCC4006 cells. BH correction-adjusted p-values were calculated, and decreases and increases in gene expression were considered significant when p-values were less than 0.05.

### Analysis of gene expression in mouse tissues

Wild-type C57BL/6 mice were treated with CDDO-Im or DMSO as a vehicle. CDDO-Im was administered at a dose of 30 μmol/kg, for which 3 mM (nmol/μl) working solution was prepared by diluting 30 mM (nmol/μl) CDDO-Im stock solution in DMSO with an isovolume of Cremophor-EL and 8x volume of PBS. The mice were sacrificed 6 hrs after the CDDO-Im administration, and representative organs were dissected. *Keap1*-knockdown (*Keap1* KD) and control wild-type mice that were obtained in the same breeding colony were sacrificed, and representative organs were dissected. The mice were used at between 7 and 9 weeks of age. Tissue samples were homogenized in ISOGEN (NIPPON GENE), and RNAs were purified. Reverse transcription and quantitative PCR were conducted in the same way as described for cell culture samples.

### Analysis of non-small cell lung cancer patients

#### Meta-analysis of the TCGA database

RNA Seq data (V2) from 517 lung adenocarcinoma tumor samples were analyzed. Tumors were considered to have high NRF2 activity if their expression levels of three or more NRF2 target genes (NQO1, GCLC, GCLM, SLC7A11, TXNRD1 and NR0B1) were ranked within the top 25%.

#### Immunohistochemistry

Rabbit polyclonal antibodies against human NOTCH3 (ab60087, Abcam) and NRF2 (sc-13032X, Santa Cruz) were used. A Histofine Kit (Nichirei Biosciences), which employs the streptavidin-biotin amplification method, was used in this study. Antigen retrieval was performed by heating slides in the microwave for 20 min (for NOTCH3) or by autoclaving slides for 5 min (for NRF2) in citric acid buffer (2 mM citric acid and 9 mM trisodium citrate dehydrate (pH 6.0)). The antigen-antibody complex was visualized using 3,3’-diaminobenzidine (DAB) solution (1 mM DAB, 50 mM Tris–HCl buffer (pH 7.6), and 0.006% H2O2) and counterstained with hematoxylin. As negative controls, we used normal rabbit IgG instead of the primary antibody or no secondary antibody. No specific immunoreactivity was detected in these sections.

#### Immunoreactivity scoring

NRF2 immunoreactivity was detected mostly in the nucleus. Cases with greater than 10% positive carcinoma cells were considered to be NRF2-positive. NOTCH3 immunoreactivity was detected in the cytoplasm and nucleus. Cases with greater than 10% positive carcinoma cells were considered to be NOTCH3-positive.

### Chromatin immunoprecipitation (ChIP) assay

ChIP was performed using anti-H3K27ac antibody (MABI0309, MAB Institute), anti-NRF2 antibody (#12721, Cell Signaling Technology), anti-FOSL2 antibody (#19967S, Cell Signaling Technology), anti-CEBPB antibody (sc-150 X, Santa Cruz) and rabbit IgG (#55944, Thermo Fisher Scientific). Cells were cross-linked with 1% formaldehyde for 10 min and lysed. The samples were sonicated to shear the DNA. Sonication was conducted according to previously described procedures (Zhang et al., 2007; Alam et al., 2017). The solubilized chromatin fraction was incubated with the primary antibodies overnight, which were prebound to anti-mouse IgG-conjugated Dynabeads (anti-H3K27ac antibody) or anti-rabbit IgG-conjugated Dynabeads (the rest of the antibodies) (Thermo Fisher Scientific). Precipitated DNA was de-crosslinked, purified and used for quantitative real-time PCR with the primers listed in Extended Table S3.

### ChIP-sequencing (ChIP-seq) analysis

ChIP was performed using control or NRF2 siRNA-treated A549 cells with an anti-H3K27ac antibody (MABI0309, MAB Institute), as described above. Precipitated DNA was de-crosslinked, purified and used for library preparation. Sequencing libraries were prepared from 1.0 or 2.0 ng of ChIPed DNA and input samples using a Mondrian SP+ system (Nugen) with an Ovation SP Ultralow DR Multiplex System (Nugen). The libraries were further purified and size-selected using an AMPure XP Kit (Beckman Coulter) and were quantified using a quantitative MiSeq (qMiSeq) method (Katsuoka et al., 2014). Optimally diluted libraries were sequenced on a HiSeq2500 (Illumina) to generate 101-base single-end reads. Sequencing files were aligned to the hg19 reference genome using Bowtie2 version 2.2.6 (Langmead and Salzberg, 2012). Reads with a mapping quality less than 20 were removed using SAMTools version 1.3.1 (Li et al., 2009). Peaks were called with MACS2 version 2.1.0.20151222 (Zhang et al., 2008). Default parameters were used for these calculations. The ENCODE blacklist (Consortium, E.P., 2012), obtained on March 22, 2016, was applied to all obtained peaks to filter out possible non-functional signals. Raw and processed data were deposited in GEO (GSE118840).

ChIP-seq data from the ENCODE project (ENCSR584GHV for NRF2, ENCSR541WQI for MAFK, ENCSR000BUB for CEBPB and the other transcription factors in A549 cells) were obtained as raw sequencing files and analyzed using the same procedure described above. Processed ChIP-seq data for H3K27ac in normal human lung samples were also obtained from the ENCODE database (ENCFF629RAA). ChIP-seq peak visualization was performed using the Integrative Genomic Viewer (Robinson et al., 2011). ENCODE ChIP-seq data for CEBPB, GR, MYC, MAX and FOSL2 were accessed through the Integrative Genomic Viewer and visualized. To clarify binding occupancy for transcription factors and H3K27ac deposition, we used *\*_treat_pileup.bdg* files generated by MACS2 with ‘--SPMR’ option. These profiles were converted into BigWig files by using KentUtils version 302 (available from https://github.com/ENCODE-DCC/kentUtils), then deepTools version 3.0.1 (Ramírez et al., 2014) was adopted to draw aggregation plots and heat maps. We used BEDtools version v2.27.1 (Quinlan, A. R., 2014) to identify overlapping between NRF2 and H3K27ac peaks.

### Correlation study of ENCODE ChIP-seq data using A549 cells

To obtain binding profiles of transcription factors in A549 cells, we analyzed ENCODE ChIP-seq data by the method described in the previous section (Extended Table S1). The experiments with an inducible treatment like ethanol or dexamethasone were excluded since specific transcriptional responses can be observed. Jaccard index values and their significance for obtained peak call results against NRF2 peaks with H3K27ac marks were calculated by GenometriCorr package version 1.1.23 (Favorov et al., 2012).

### Production of virus particles

For lentiviral infection, lentiviral and packaging vectors were transfected into 293FT cells. For retroviral infection, retroviral vectors were transfected into PLAT-A cells. The culture media were replaced with fresh media 24 hrs after transfection. The cells were incubated for an additional 24 hrs, and the culture supernatants were used as the lentivirus or retrovirus particle sources.

### Disruption of *NOTCH3* enhancer to establish ΔN3U cells

Two guide RNAs (gRNAs) were designed to disrupt NRF2 binding sites (antioxidant responsive elements; AREs) located 10-kb upstream of the transcription start site of the *NOTCH3* gene. Lentiviral vectors expressing these gRNAs together with Cas9 mRNA were constructed by inserting annealed oligoDNAs (Extended Table S4) into LentiCRISPRv2 (Addgene). H460, A549 and H2023 cells were infected with lentiviral particles with 12.5 μg/ml polybrene. After incubation for 24 hrs, the cells were re-plated in 10 cm dishes and incubated in selection medium containing 2 μg/ml puromycin. Single clones were selected using cloning rings (TOHO). DNA was purified from each clone, and the modified regions were amplified using the primer sets listed in Extended Table S5. The PCR products were cloned and sequenced to verify disruption of the NRF2 binding sites.

### Establishment of inducible *NRF2* knockdown cells

The inducible *NRF2* shRNA lentiviral vector (SMARTvector Inducible Lentiviral shRNA; V3SH11252, V3IHSMCG_6358637 and V3IHSMCG_6804335) and control vector (SMARTvector Inducible Non-targeting mCMV-TurboGFP; VSC11651) were purchased from Dharmacon. H460 cells were infected with lentiviral particles with 12.5 μg/ml polybrene. After 24 hrs, the cells were re-plated in 10 cm dishes and cultured in selection medium containing 2 μg/ml puromycin. 10 μg/ml tetracycline (APOLLO) was added 48 hrs prior to initiation of the experiments using these cell lines.

### Introduction of LacZ and NOTCH3 intracellular domain (N3ICD) into ΔN3U cells

Expression vectors for N3ICD (pLV[Exp]-CMV>{hNotch3ICD}:IRES:EGFP(ns):T2A:Bsd) and LacZ (pLV[Exp]-CMV>LacZ:IRES:EGFP(ns):T2A:Bsd) were purchased from Vector Builder. ΔN3U H460 cells were infected with lentiviral particles with 12.5 μg/ml polybrene. After incubation for 24 hrs, the cells were re-plated in 10 cm dishes and incubated in selection medium containing 2-5 μg/ml blasticidin.

### Spheroid formation assay

Cell growth was examined in spheroid culture. 10^3^ cells with 100 μl culture media were seeded in each well of a low-attachment U-bottom plate (PrimeSurface 96 well Plate, MS-9096UZ, Sumitomo Bakelite Co.) followed by transfection with 2 pmol/well of *NRF2* siRNA (HSS107128) or control siRNA using RNAiMAX (Invitrogen). Spheroids were observed 24 and 96 hrs after transfection. After 96 hrs, cell proliferation was assessed using the Cell Counting Kit-8 (Nacalai Tesque) according to the manufacturer’s protocol.

### Oncosphere formation assay

Oncospheres were cultured as previously described (Justilien et al., 2012). Briefly, cells were cultured in ultra-low attachment dishes (Corning) in CSC medium. The CSC medium consisted of serum-free DMEM-F12 medium (Gibco-Invitrogen) containing 50 μg/ml insulin (Sigma-Aldrich), 0.4% Albumin Bovine Fraction V (Sigma-Aldrich), N-2 Plus Media Supplement (R&D Systems), Gibco B-27 Supplement (Thermo Fisher Scientific), 20 ng/ml EGF (Pepro Tech) and 10 ng/ml bFGF (Pepro Tech). A549 (1 x 10^4^ cells), H2023 (4 x 10^4^ cells) and H460 (1 x 10^4^ cells) cells were seeded in each well in 2 ml CSC medium. Cells were harvested 7 days after seeding. Transfections with control siRNA and siRNAs against *NRF2* and *CEBPB* were conducted concurrently with cell seeding in CSC medium, and culture media were left unchanged until the cell harvest. Viable cells were counted using trypan blue staining. Whole cell proteins were prepared for immunoblotting.

### Xenograft experiments

H460, A549 and H2023 cell suspensions were combined with Matrigel (Corning) and subcutaneously injected into the flank of four-week-old male Balbc nu/nu mice. In the case of the inducible *NRF2*-knockdown H460 cells, 1 x 10^4^ cells were injected and the resulting tumors were analyzed after 17 days. In the case of the *NOTCH3* enhancer-disrupted (ΔN3E) H460 cells, 1 x 10^3^ and 1 x 10^4^ cells were injected and the resulting tumors were dissected and weighed after 21 days and 15 days, respectively. In the case of ΔN3E A549 and H2023 cells, 1 x 10^5^ (A549) and 3 x 10^5^ (H2023) cells were injected and both of the resulting tumors were dissected and weighed after 35 days. In the case of ΔN3E H460 cells with LacZ expression and those with N3ICD expression, 1 x 10^4^ cells were injected and the resulting tumors were analyzed after 18 days. In the CDDP administration experiments, 0.3mg/kg of CDDP or normal saline as a vehicle were injected into the peritoneal cavity of the Balbc nu/nu mice once a week starting from the timing of H460 cell transplantation.

### Serial transplantation experiments

Tumors from the primary xenograft experiment were dissected from mice and chopped into small pieces under sterile conditions and incubated at 37°C for 1 hr in 5 ml low glucose DMEM supplemented with 10% fetal bovine serum (FBS) and penicillin/streptomycin containing 1 mg/mL collagenase type IV (Sigma-Aldrich, #C5138) and 100 μg/ml DNase I (SIGMA). The samples were incubated with 5 ml RBC lysis buffer (155 mM NH_4_Cl, 15 mM NaHCO_3_, 0.1 M EDTA, pH7.3) for 10 min, followed by filtration to remove debris. The samples were then incubated with anti-biotinylated CD31 (#13-0311-81, eBioscience) and CD45 (#13-0451-85, eBioscience) antibodies, followed by reaction with Dynabeads M-280 streptavidin (Thermo Fisher Scientific) to remove murine cells of endothelial and hematopoietic origins. To further remove the remaining murine cells, the samples were reacted with anti-mouse MCH class I antibody (ab95572, Abcam), and human tumor cells were collected as the mouse MCH class I-negative fraction using flow cytometry (BD FACSAria, Becton Dickinson). Then, 1 x 10^3^ WT and ΔN3E H460 cells were used for the secondary xenograft experiment. The time course of the experiment is shown in Extended Figure 13a.

### Immunoprecipitation

A549 cells were harvested using a cell scraper and washed three times in 1 x PBS. The cell pellet was resuspended in 1 x PBS containing 0.5 mM DTME and 0.5 mM DSP and incubated at room temperature for 30 min, followed by incubation in quenching buffer (20 mM Tris-HCl (pH 7.5), 5 mM cysteine) at 25°C for 5 min. After washing in ice-cold PBS, the pellet was sonicated in RIPA buffer briefly and centrifuged at 14,000 g at 4°C for 5 min. The supernatant was subjected to anti-NRF2 affinity purification. An anti-NRF2 antibody (#12721, Cell Signaling Technology) and control rabbit IgG (#55944, Thermo Fisher Scientific) were crosslinked to a 1:1 mixture of Dynabeads protein A and protein G (Thermo Fisher Scientific) with DMP and incubated with the supernatant at 4°C for 2 hr. After washing in RIPA buffer, the NRF2 complex was eluted from the beads by incubation in elution buffer (50 mM Tris-HCl (pH 8.0), 0.2 M NaCl, 2w/v% SDS, 50 mM DTT) at 37°C for 30 min. The eluate was analyzed by immunoblot analysis with anti-NRF2 antibody (sc-13032X, Santa Cruz), anti-FOSL2 antibody (#19967S, Cell Signaling Technology), and anti-CEBPB antibody (sc-150 X, Santa Cruz).

### Quantification and statistical analysis

Statistical significance was evaluated using an unpaired two-sample Student’s *t*-test, the Wilcoxon rank sum test and one-way ANOVA followed by the Bonferroni post hoc test. Confidence intervals were calculated for all fold change evaluations. Associations between NRF2- and NOTCH3-staining were evaluated by a cross-table using a chi-square test. Kaplan-Meier analysis was performed for cumulative and relapse-free survival, and statistical significance was evaluated using the log-rank test. Univariate and multivariate analyses were evaluated using the Cox proportional hazards model. These analyses were performed using Microsoft Office Excel (Microsoft), Prism 7 (GraphPad Software, Inc.), JMP Pro 13 and StatView 5.0J software (SAS Institute). *P*<0.05 and α<0.05 (confidence interval) were considered to be statistically significant.

### Data and code availability

RNA-seq data generated in this study were deposited in GEO (GSE118840, GSE118842). ChIP-seq data generated in this study were deposited in GEO (GSE118840).

## Author Contributions

K.O., H.A., Z.L., N.O., H.K., Y. Onodera, M.A.A., D.M., Takuma S., F.K., and N.O. conducted the experiments, analyzed the data and wrote the paper. S.T. and I.M. analyzed the data. M.W., A.S., Y. Okada and Takashi S. analyzed the clinical data. M.Y. provided critical biomaterials and an analysis platform for the study and analyzed the data. K.K. supervised the research, analyzed data and wrote the paper. H.S. conducted the experiments, supervised the research, analyzed the data and wrote the paper. H.M. designed the study, supervised the research, analyzed the data and wrote the paper.

## Acknowledgments

We thank Dr. Christian Siebel for critical reading of the manuscript, Dr. Hideyuki Saya for advice on cancer stemness and Drs. Kazuhiko Igarashi, Hiroki Shima and Kyoko Ochiai for advice on NRF2 complex purification. We also thank Ms. Nozomi Hatanaka for deep sequencing technical support and the Biomedical Research Core of the Tohoku University Graduate School of Medicine for their technical support.

This work was supported by JSPS under grant numbers 18H02621 (HM), 18H04794 (HM), 17F17116 (HM), 17K08618 (HS) and 16K12519 (KK), the Naito Foundation (HM), a research grant from the Princess Takamatsu Cancer Research Fund 15-24728 (HM), the Uehara Memorial Foundation (HM), a research grant from the Gonryo Medical Foundation (HS) and AMED under grant numbers JP18am0101067 (KK) and JP19gm5010002 (HM). The funders had no role in the study design, data collection and analysis, decision to publish or manuscript preparation.

## Notes

### Competing Interest Statement

The authors have declared no competing interest.

