## Extended Figure 1-14 for "Enhancer Remodeling Promotes Tumor-Initiating Activity in NRF2-Activated Non-Small Cell Lung Cancers"

Short title: NRF2-NOTCH3 axis promotes tumor-initiating activity of cancer

***Corresponding authors**

Hiroki Sekine, Ph.D.,

Department of Gene Expression Regulation, Institute of Development, Aging and Cancer, Tohoku University.

4-1 Seiryo-cho, Aoba-ku, Sendai, Miyagi, 980-8575, Japan.

.

Hozumi Motohashi, M.D., Ph.D.,

Department of Gene Expression Regulation, Institute of Development, Aging and Cancer, Tohoku University.

4-1 Seiryo-cho, Aoba-ku, Sendai, Miyagi, 980-8575, Japan.

.

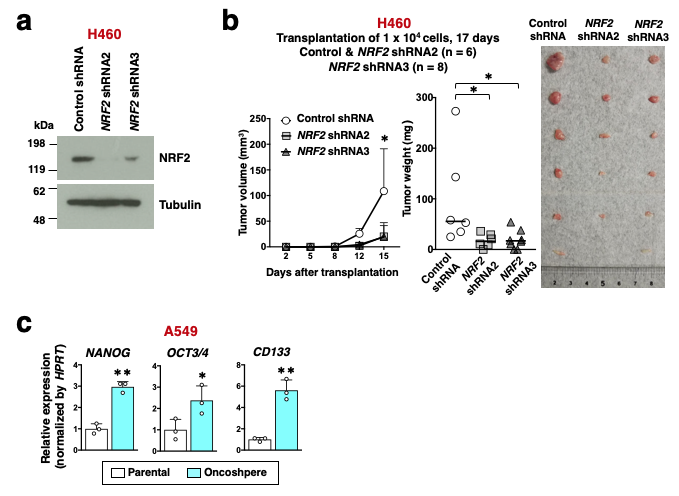

**Extended Figure 1. Evaluation of the tumor-initiating activity of NRF2-activated NSCLCs.**

**a.** Immunoblot analysis of NRF2 protein levels in inducible *NRF2*-knockdown H460 cells (*NRF2* shRNA2 and *NRF2* shRNA3) and control H460 cells after 48 hrs of DOX treatment. Two different shRNAs were used for inducible knockdown of *NRF2*. Tubulin was used as the loading control. The results shown are representative of three independent experiments. **b.** Xenograft experiment using inducible *NRF2*-knockdown and control H460 cells. 1 x 10^4^ cells were mixed with Matrigel and subcutaneously transplanted into nude mice. Tumors were weighed after 17 days. A photograph shows xenograft tumors at the time of tumor weight measurement. Horizontal bars indicate median levels. The Wilcoxon rank sum test was performed. **p*<0.05. **c.** RT-PCR measuring the expression of stem cell marker genes normalized to *HPRT* in parental and oncosphere-forming A549 cells. The average and SD of three independent experiments are shown. The Student’s *t* test was performed. **p*<0.05, ***p*<0.01.

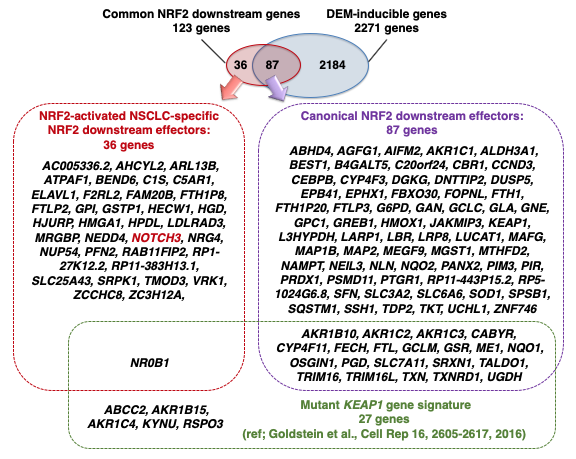

**Extended Figure 2. NRF2 downstream effectors identified in RNA-seq analysis with NRF2-activated and NRF2-normal NSCLC cell lines.**

87 canonical NRF2 downstream effectors and 36 NRF2-activated cancer-specific NRF2 downstream effectors were compared with the mutant *KEAP1* gene signature (Goldstein et al., 2016).

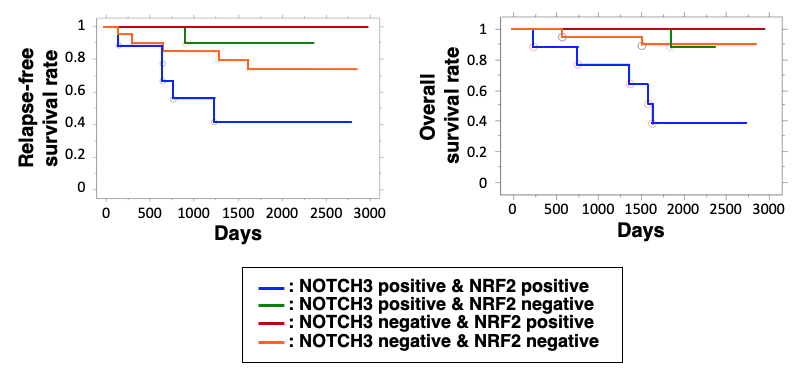

**Extended Figure 3. Prognosis of lung adenocarcinoma patients according to NRF2 and NOTCH3 status.**

Overall survival rates and relapse-free survival rates of lung adenocarcinoma patients according to NRF2 and NOTCH3 status.

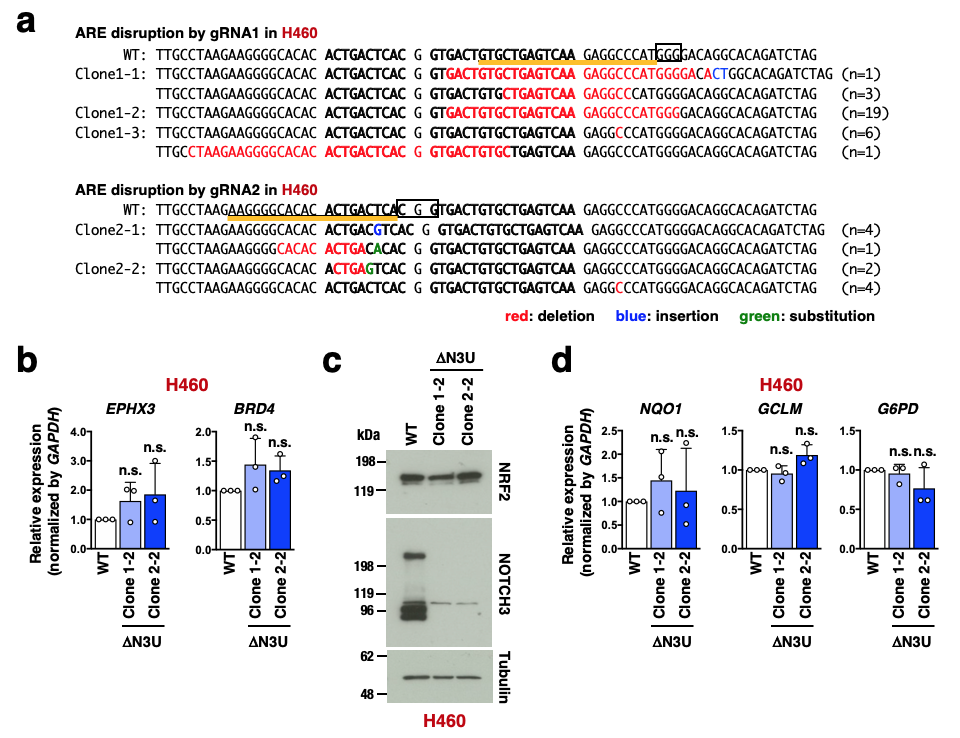

**Extended Figure 4. Genome editing of the *NOTCH3* upstream region in H460 cells.**

**a.** DNA sequences of NRF2-binding regions were altered by CRISPR-Cas9 genome editing in the *NOTCH3* enhancer of H460. Sequences corresponding to gRNAs are underlined in orange, and protospacer-adjacent motifs are boxed. PCR products were cloned and sequenced using primer pairs flanking the targeted regions. Deleted, inserted and substituted bases are indicated in red, blue and green, respectively. Numbers of altered sequences obtained in the cloned PCR products are shown to the right of the figure. **b, d.** RT-PCR measuring the expression of genes neighboring *NOTCH3* (**b**) and canonical NRF2 target genes (**d**) normalized to *GAPDH* in ΔN3U and WT H460 cells. Fold changes of the normalized values were calculated in comparison to WT H460 cells. The average and SD of the fold changes from three independent experiments are shown. Confidence interval estimation was conducted for the ΔN3U H460 clones. n.s.: not significant. **c.** Immunoblot analysis of NOTCH3 and NRF2 protein expression. Tubulin expression was used as a loading control. The results shown are representative of three independent experiments.

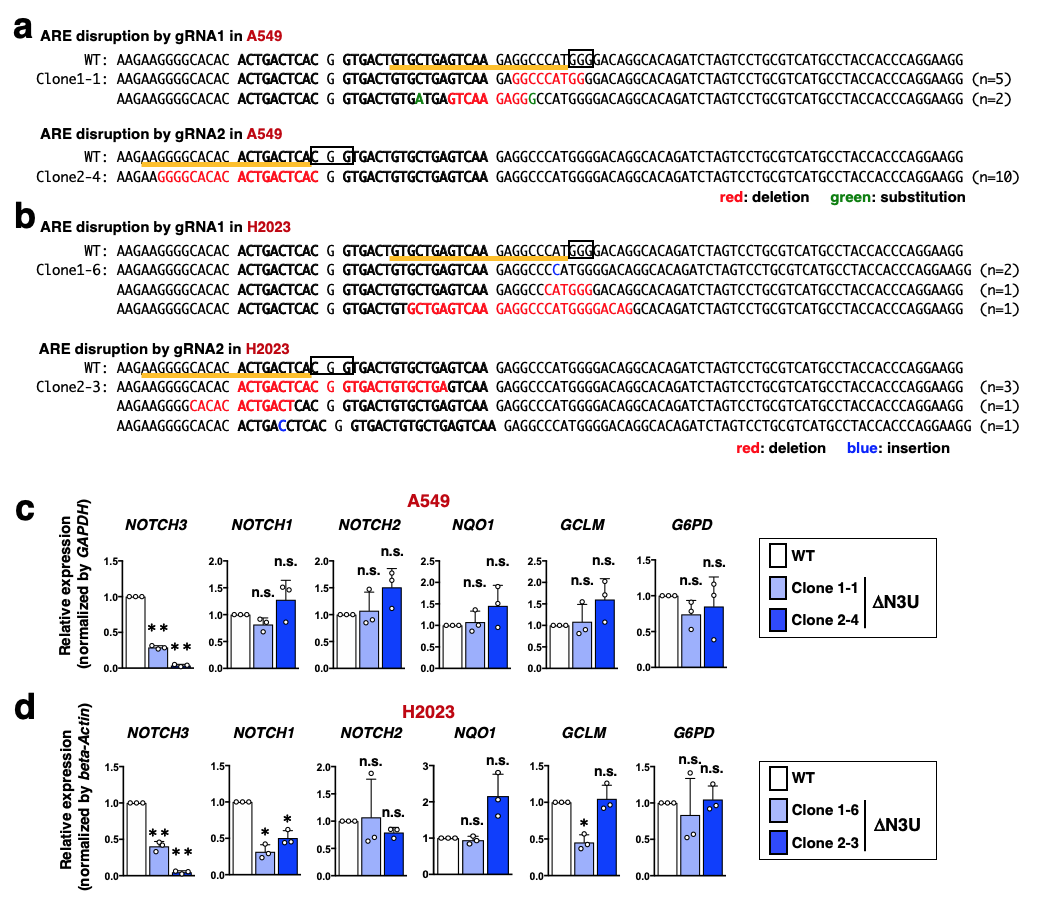

**Extended Figure 5. Genome editing of the *NOTCH3* upstream region in A549 and H2023 cells.**

**a, b.** DNA sequences of NRF2-binding regions were altered by CRISPR-Cas9 genome editing in the *NOTCH3* enhancer of A549 (**a**) and H2023 (**b**) cells. Sequences corresponding to gRNAs are underlined in orange, and protospacer-adjacent motifs are boxed. PCR products were cloned and sequenced using primer pairs flanking the targeted regions. Deleted, inserted and substituted bases are indicated in red, blue and green, respectively. Numbers of altered sequences obtained in the cloned PCR products are shown to the right of the figure. **c, d.** RT-PCR measuring the expression of the *NOTCH* family and canonical NRF2 target genes in ΔN3U and WT A549 cells normalized to *GAPDH* (**c**) and ΔN3U and WT H2023 cells normalized to *beta-Actin* (**d**). Fold changes of the normalized values were calculated in comparison to WT A549 and WT H2023 cells. The average and SD of the fold changes from three independent experiments are shown. Confidence interval estimations were conducted for the ΔN3U A549 and ΔN3U H2023 clones. **a*<0.05, ***a*<0.01, n.s.: not significant.

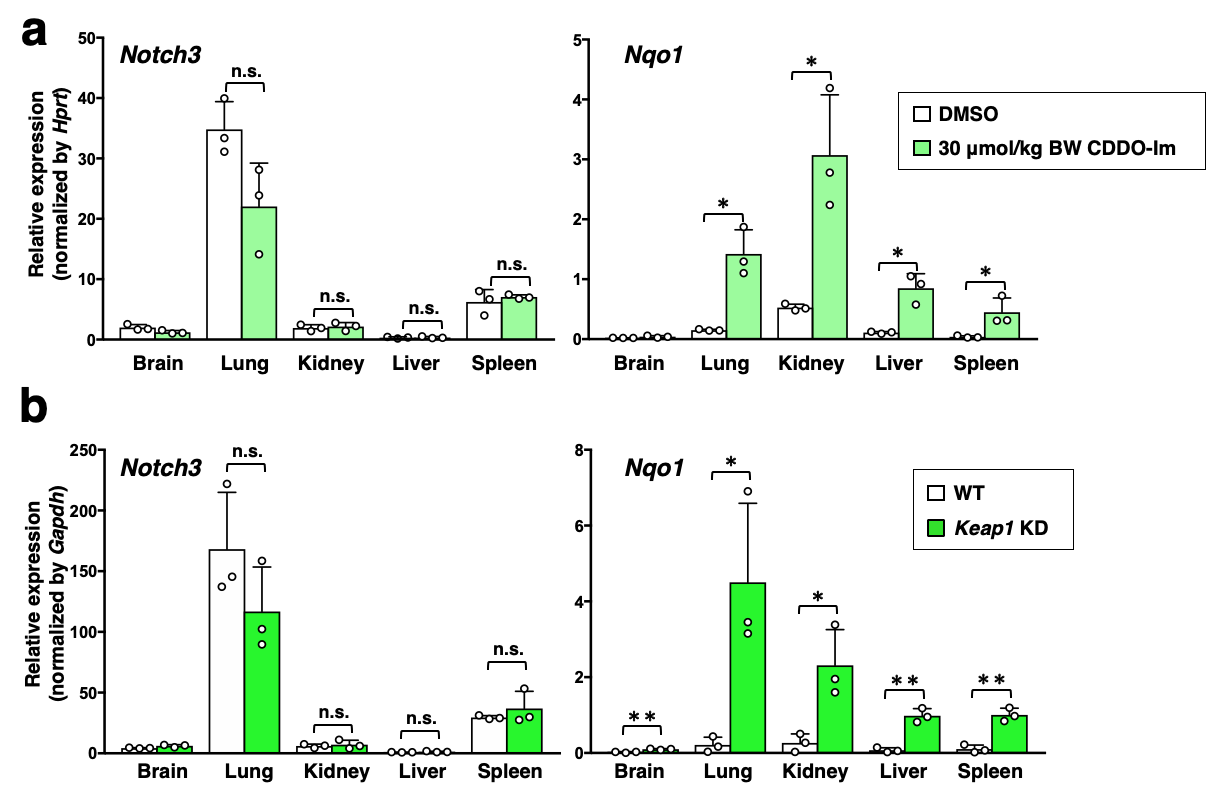

**Extended Figure 6. *Notch3* expression in normal mouse tissues.**

**a.** RT-PCR measuring the expression of *Notch3* and *Nqo1* normalized to *Hprt* in representative tissues of mice treated with vehicle (DMSO) or CDDO-Im. The average and SD of three independent experiments are shown. The Student’s *t* test was performed. **p*<0.05, n.s.: not significant. **b.** RT-PCR measuring the expression of *Notch3* and *Nqo1* normalized to *Gapdh* in representative tissues of WT and *Keap1* knockdown mice. The average and SD of three independent experiments are shown. The Student’s *t* test was performed. **p*<0.05, ***p*<0.005, n.s.: not significant.

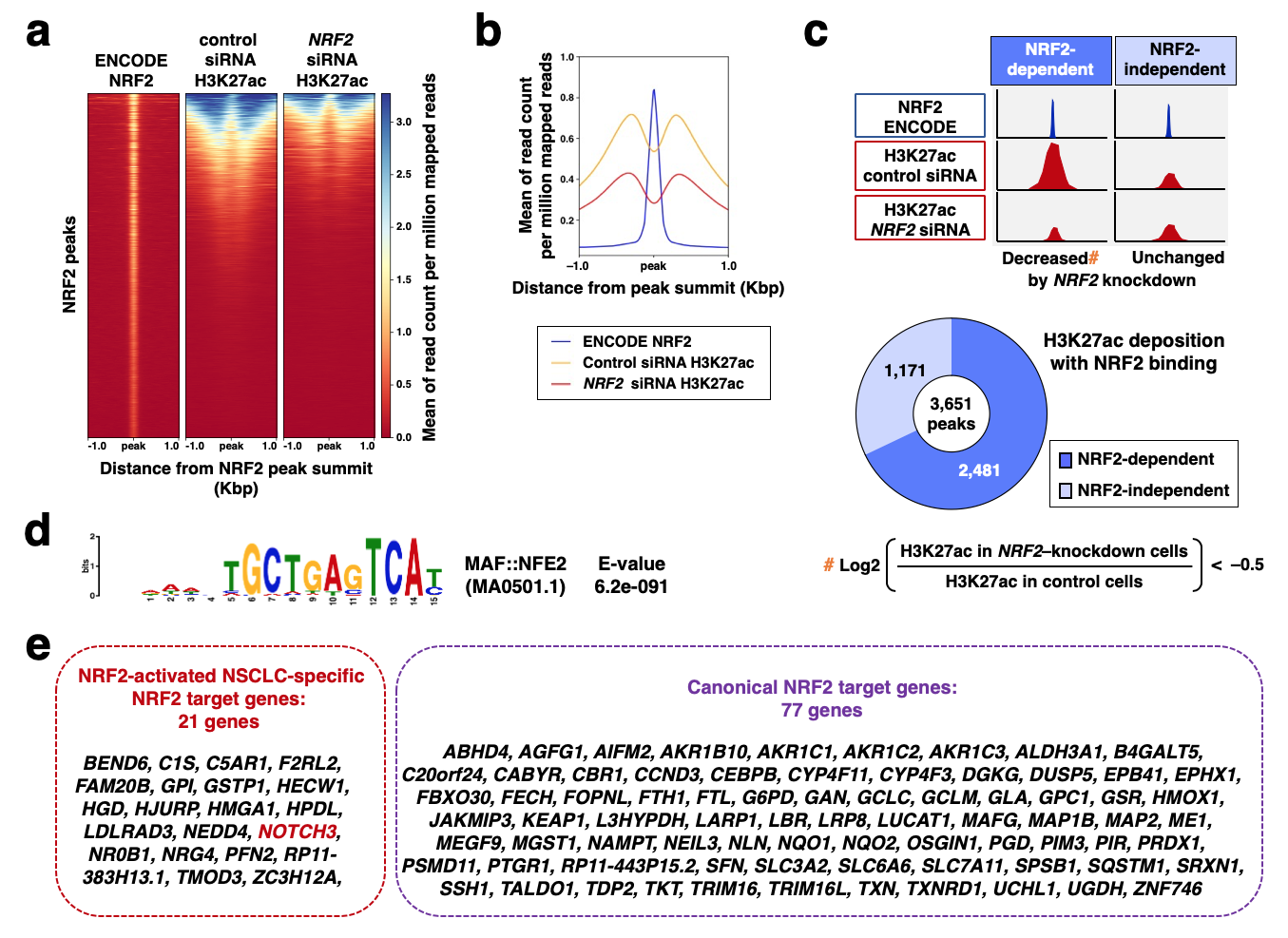

**Extended Figure 7. NRF2 contributes to unique enhancer formation in NRF2-activated NSCLC cells.**

**a, b.** ChIP-seq analysis using an antibody against H3K27ac in A549 cells treated with control siRNA or *NRF2* siRNA. A heat map (**a**) and an aggregation plot (**b**) show acetylated H3K27 (H3K27ac) deposition surrounding NRF2 binding sites obtained from the ENCODE database. **c.** Classification of NRF2-bound enhancers according to the NRF2 dependency of H3K27ac deposition. **d.** A motif enriched in NRF2-dependent enhancers obtained using the MEME-ChIP platform. **e.** NRF2-activated NSCLC-specific NRF2 target genes and canonical NRF2 target genes based on the presence of NRF2-dependent enhancers. **e.** NRF2-activated NSCLC-specific NRF2 target genes and canonical NRF2 target genes based on the presence of NRF2-dependent enhancers.

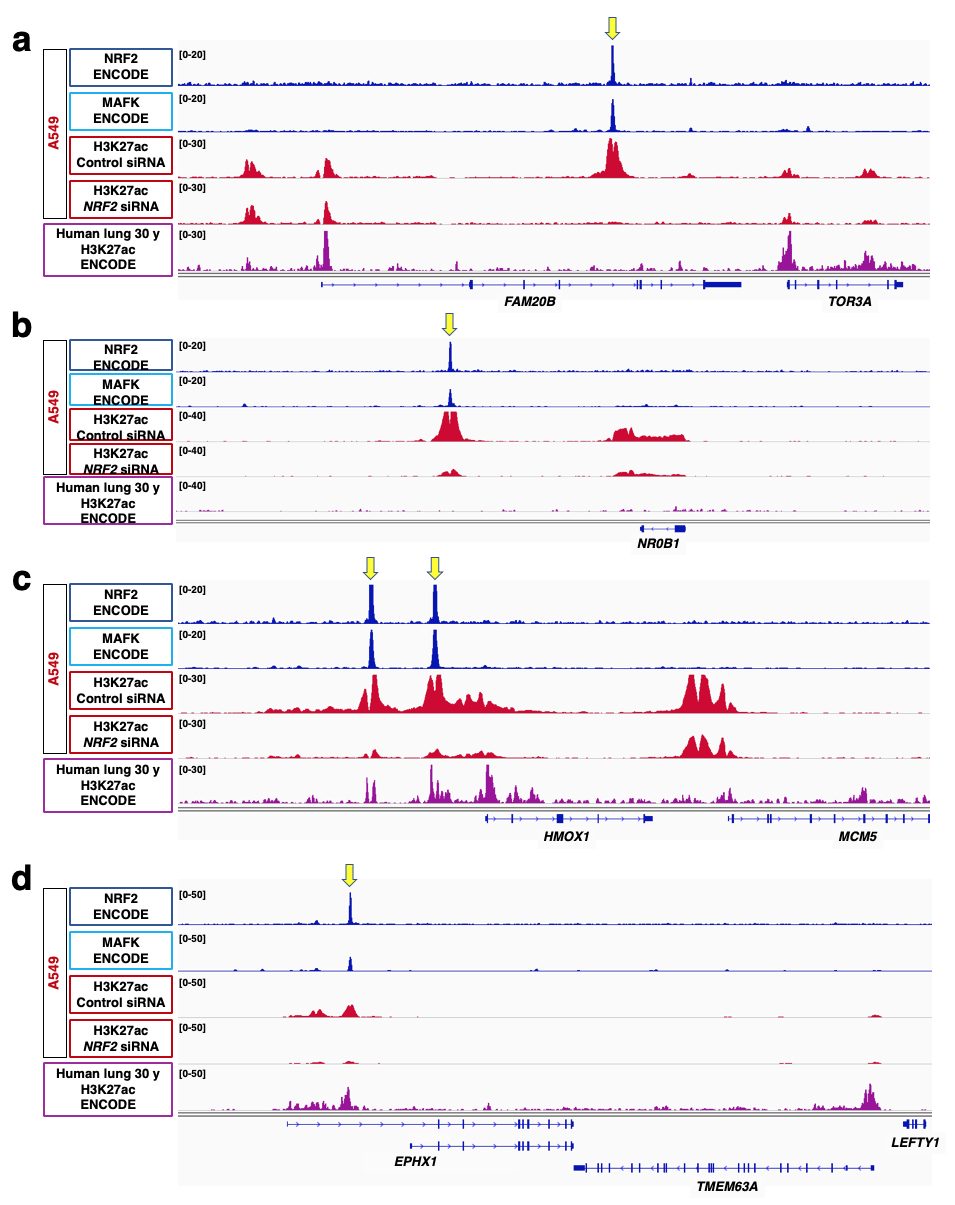

**Extended Figure 8. ChIP-seq profiles at representative NRF2-activated NSCLC-specific and canonical NRF2 target loci.**

Genome browser views of NRF2-activated NSCLC-specific NRF2 target loci (**a**, **b**) and canonical NRF2 target loci (**c**, **d**). A549 cells (upper panels) and normal human lung samples (lower panel) are shown. NRF2 and MAFK chromatin binding in A549 cells and H3K27ac deposition patterns in normal human lung samples were obtained from the ENCODE database. Acetylated H3K27 deposition profiles in A549 cells treated with control siRNA or *NRF2* siRNA were obtained in this study. Yellow arrows indicate NRF2 binding peaks in the respective NRF2 target loci.

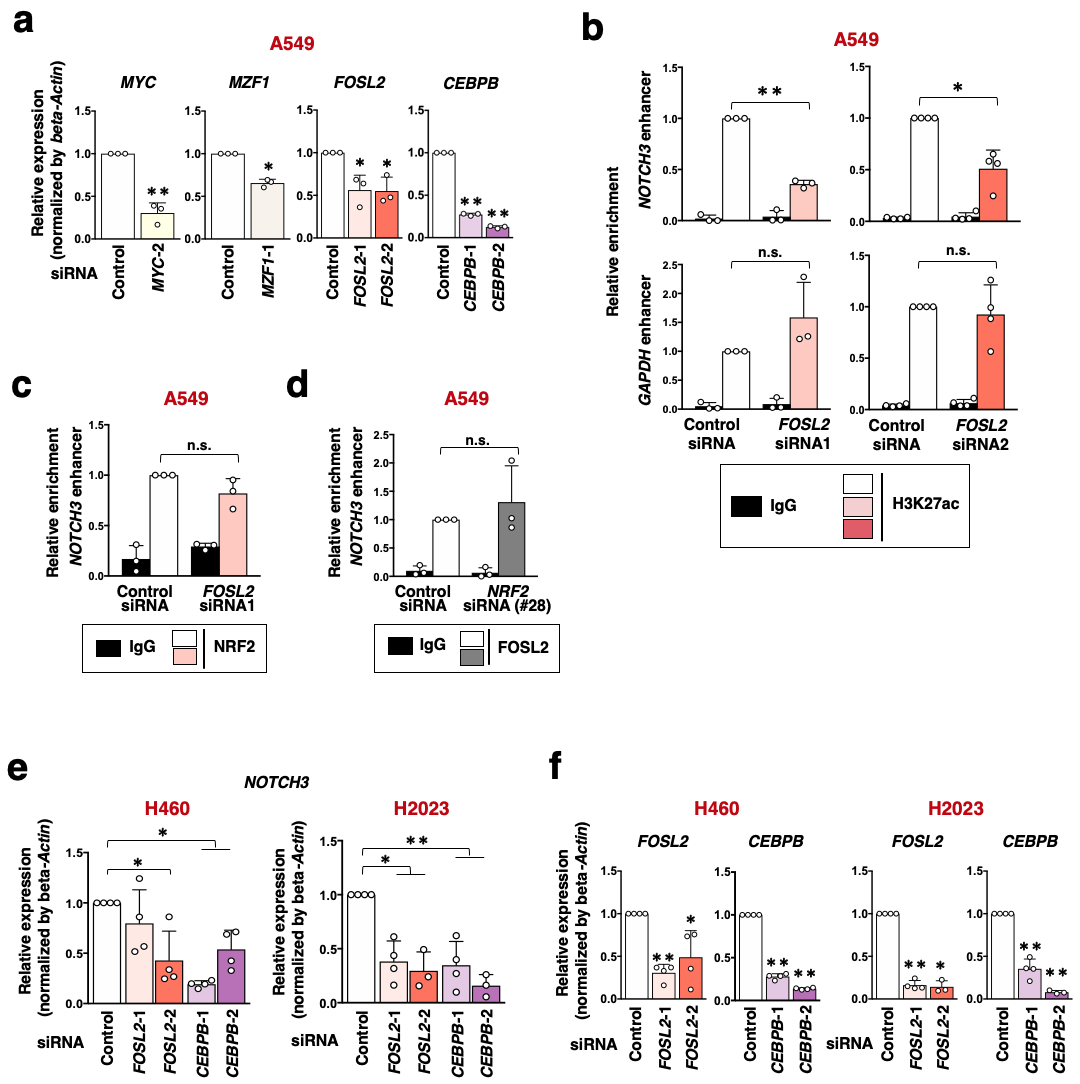

**Extended Figure 9. CEBPB cooperates with NRF2 and contributes to *NOTCH3* enhancer formation in NRF2-activated NSCLCs.**

**a.** RT-PCR measuring the knockdown efficiency of siRNAs against *MYC*, *MZF1*, *FOSL2* and *CEBPB* in A549 cells. Fold changes of the normalized values were calculated in comparison to control siRNA-treated cells. The average and SD of the fold changes from three independent experiments are shown. Confidence interval estimation was conducted to evaluate statistical significance. **a*<0.05, ***a*<0.01. **b, c.** ChIP assay using H3K27ac (**b**) and NRF2 (**c**) antibodies in A549 cells treated with *FOSL2* or control siRNAs. Enrichment of the *NOTCH3* enhancer region was examined (**b**, **c**). The *GAPDH* enhancer was selected as a control locus (**b**). Fold changes of %input values were calculated in comparison to the control cells reacted with H3K27ac or NRF2 antibody. The average and SD of 3 or 4 independent experiments are shown. Confidence interval estimation was conducted for knockdown samples reacted with H3K27ac or NRF2 antibody. **a*<0.05, n.s.: not significant. **d.** ChIP assay using FOSL2 antibody in A549 cells treated with NRF2 or control siRNAs. Enrichment of the *NOTCH3* enhancer region was examined. Fold changes of %input values were calculated in comparison to A549 cells with control siRNA. The average and SD of three independent experiments are shown. Confidence interval estimation was conducted for A549 cells with *NRF2* siRNA. ***a*<0.01, n.s.: not significant. **e.** RT-PCR measuring the expression of *NOTCH3* normalized to *beta-Actin* in H460 and H2023 cells treated with *FOSL2*, *CEBPB* and control siRNAs. Fold changes of the normalized values were calculated in comparison to control siRNA-treated cells. The average and SD of the fold changes from four independent experiments are shown. Confidence interval estimation was conducted to evaluate statistical significance. **a*<0.05, ***a*<0.01. **f.** RT-PCR measuring the knockdown efficiency of siRNAs against *FOSL2* and *CEBPB* in H460 and H2023 cells. Fold changes of the normalized values were calculated in comparison to control siRNA-treated cells. The average and SD of the fold changes from four independent experiments are shown. Confidence interval estimation was conducted to evaluate statistical significance. **a*<0.05, ***a*<0.01.

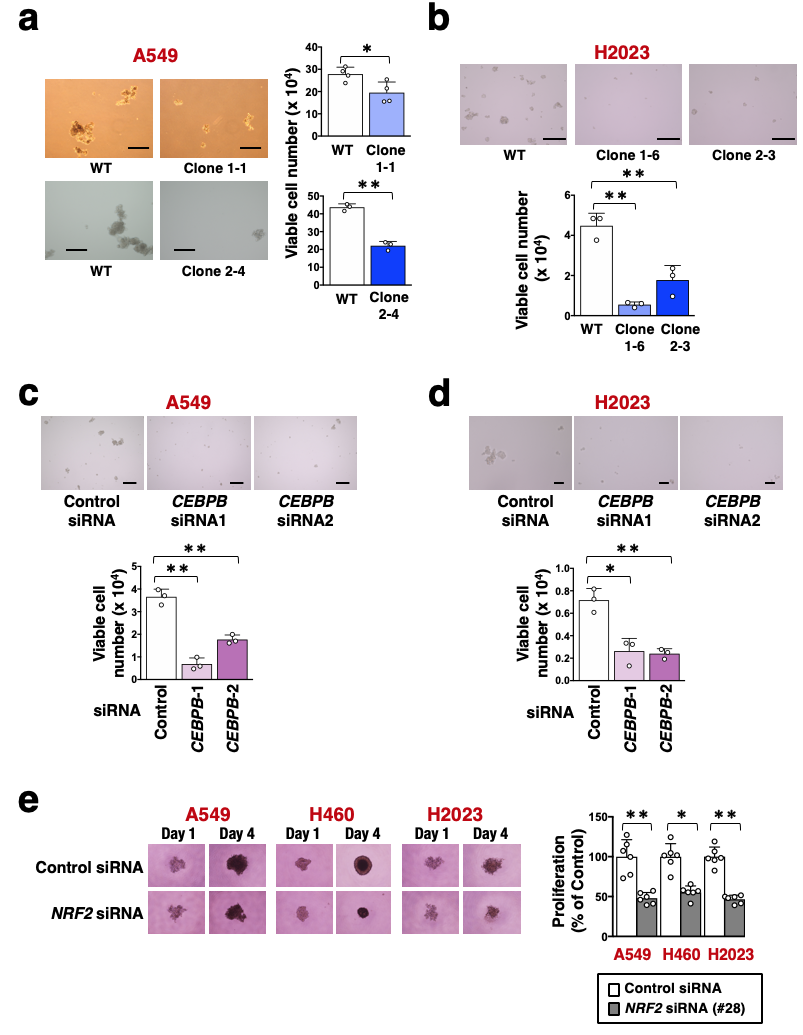

**Extended Figure 10. *NOTCH3* enhancer promotes oncosphere growth of NRF2-activated NSCLC cell lines.**

**a.** Oncosphere growth of ΔN3U and WT A549 cells (left panels). Scale bars indicate 500 μm. Viable cells were counted after trypsinization (right panels). Average cell numbers and SD from three or four independent experiments are shown. The Student’s *t* test was performed. **p*<0.05, ***p*<0.01. **b.** Oncosphere growth of ΔN3U and WT H2023 cells (upper panels). Scale bars indicate 300 μm. Viable cells were counted after trypsinization (lower panel). Average cell numbers and SD from three independent experiments are shown. The Student’s *t* test was performed. ***p*<0.01. **c, d.** Oncosphere growth of A549 (**c**) and H2023 (**d**) with or without *CEBPB* knockdown. Scale bars indicate 300 μm (**c**) and 100 μm (**d**). Viable cells were counted after trypsinization (right panels). Average cell numbers and SD from three independent experiments are shown. The Student’s *t* test was performed. **p*<0.05, ***p*<0.01. **e.** Spheroid growth of A549, H2023 and H460 cells. Spheroids are shown at 1 and 4 days after transfection of *NRF2* siRNA (upper panels). Cell numbers were estimated using a cell counting kit on day 4 (lower panel). Average cell numbers and SD from six independent experiments are shown. The average cell numbers from NSCLC cell lines treated with control siRNA were set as 100%. The Student’s *t* test was performed. **p*<0.05, ***p*<0.01.

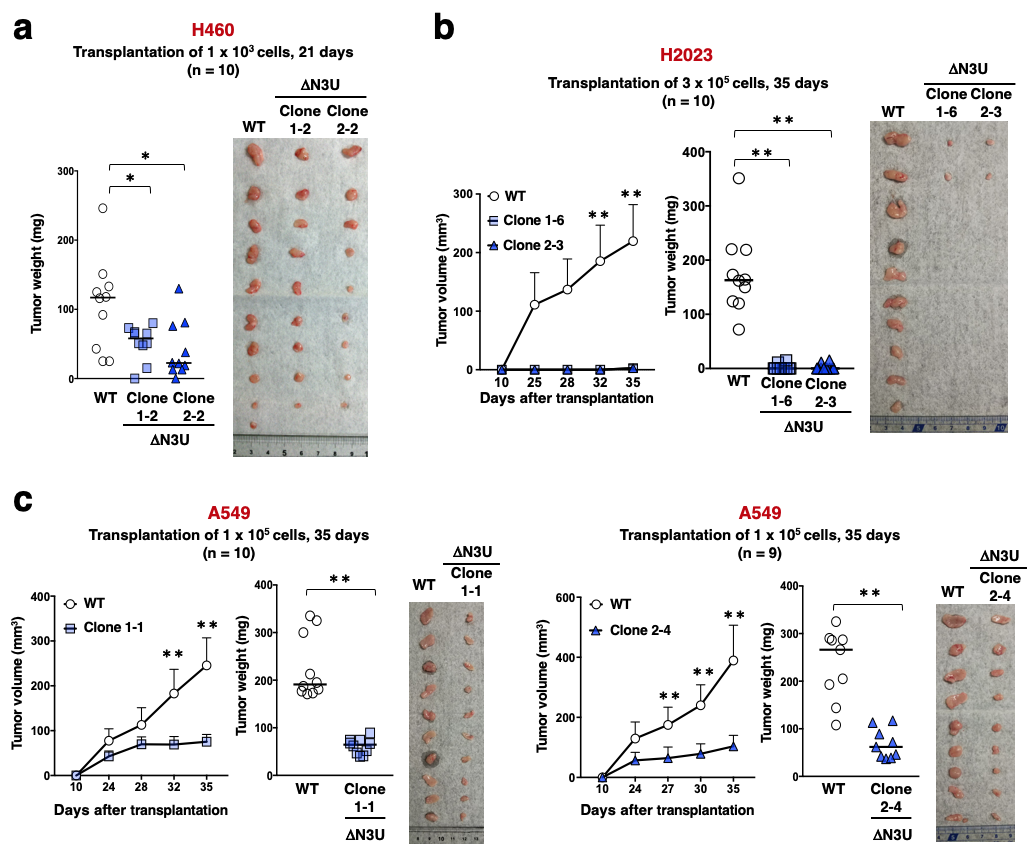

**Extended Figure 11. *NOTCH3* enhancer promotes tumor-initiating activity of NRF2-activated NSCLCs.**

**a.** Xenograft experiment using ΔN3U and WT H460 cells. 1 x 10^3^ cells were mixed with Matrigel and subcutaneously transplanted into nude mice. Tumors were weighed after 21 days. A photograph shows xenograft tumors at the time of tumor weight measurement. Horizontal bars indicate the median tumor weight. The Wilcoxon rank sum test was performed. **p*<0.05. **b, c.** Xenograft experiments using ΔN3U and WT H2023 (**b**) and A549 (**c**) cells. 3 x 10^5^ (**b**) and 1 x 10^5^ cells (**c**) were mixed with Matrigel and subcutaneously transplanted into nude mice. Tumors were weighed after 35 days in both cells. Photographs show xenograft tumors at the time of tumor weight measurement. Horizontal bars indicate the median tumor weight. The Wilcoxon rank sum test was performed. ***p*<0.005.

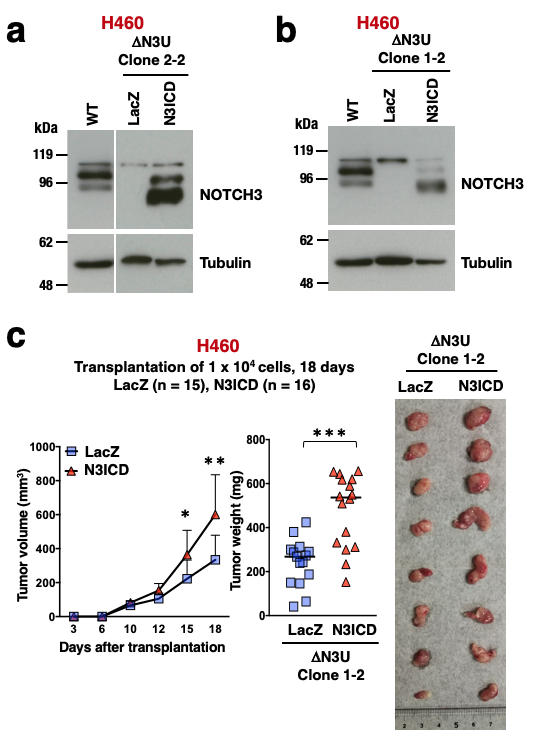

**Extended Figure 12. NOTCH3 intracellular domain restores the tumorigenesis of ΔN3U H460 cells.**

**a,b.** Immunoblot analysis of NOTCH3 protein levels of WT H460 cells and ΔN3U H460 Clone 2-2 cells (**a**), Clone 1-2 cells (**b**) with LacZ and N3ICD expression. Tubulin expression was used as a loading control. The results shown are representative of three independent experiments. **c.** Xenograft experiment using ΔN3U H460 Clone 1-2 cells expressing LacZ and N3ICD. 1 x 10^4^ cells of Clone 1-2 were mixed with Matrigel and subcutaneously transplanted into nude mice. Tumors were weighed after 18 days. A photograph shows representative xenograft tumors at the time of tumor weight measurement. Horizontal bars indicate the median tumor weight. The Wilcoxon rank sum test was performed. **p*<0.05, ***p*<0.005, ****p*<0.0005.

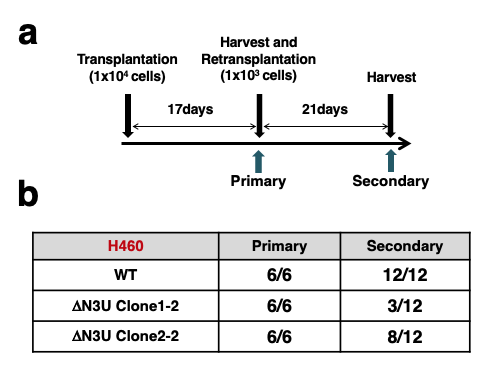

**Extended Figure 13. Serial transplantation experiment of H460 and its mutant cells.**

**a.** Protocol for the serial transplantation experiment. **b.** Tumor numbers in the primary and secondary transplantation.

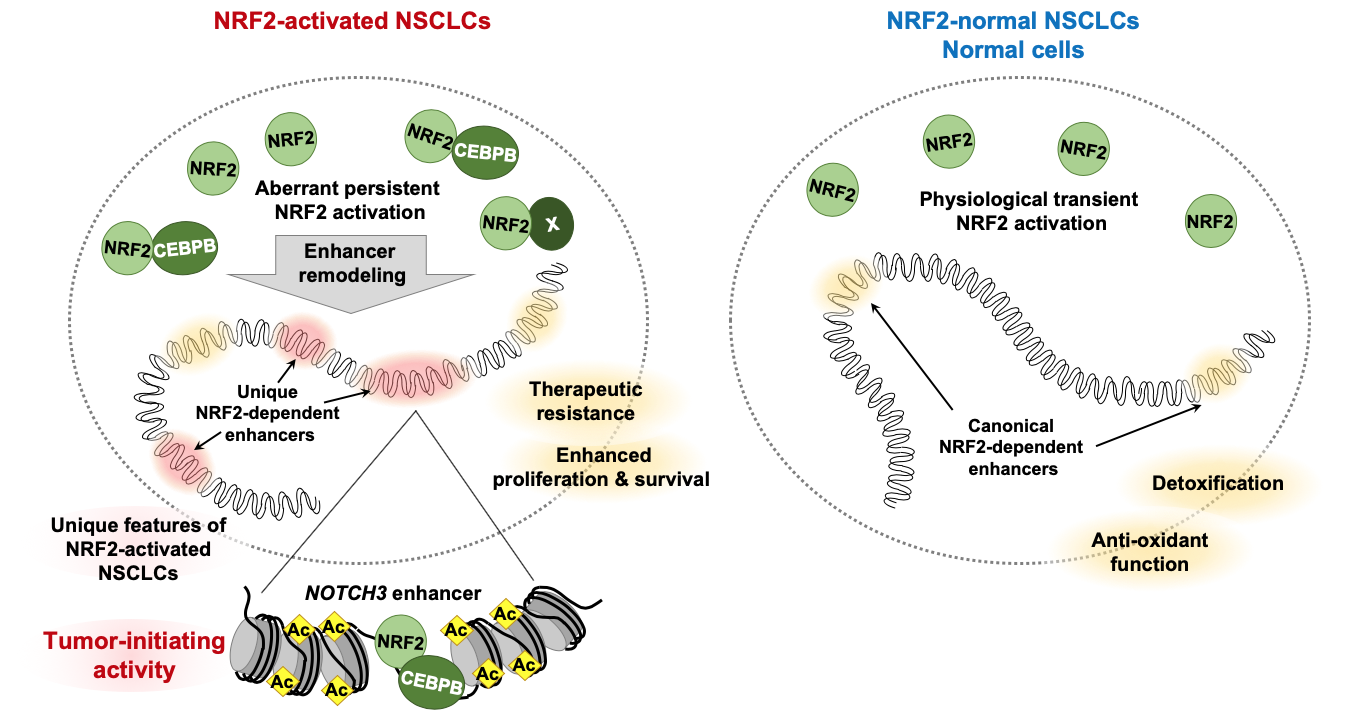

**Extended Figure 14. Illustration of enhancer remodeling in NRF2-activated NSCLCs.**

Under physiological conditions and in NRF2-normal NSCLCs, NRF2 is transiently activated in response to stimuli and induces canonical enhancer formation to activate genes involved in detoxification and antioxidant function (right). The canonical enhancers are regarded to confer therapeutic resistance and promote cell proliferation and survival. In NRF2-activated NSCLCs, enhancer remodeling creates unique NRF2-dependent enhancers (left). CEBPB is one of the cooperative factors for NRF2 that mediates enhancer remodeling. The *NOTCH3* enhancer, one of the unique enhancers, promotes tumor-initiating activity.
